# Next generation protein-corrole bio-assemblies provide effective tumoricidal treatment in a metastatic triple-negative breast cancer model

**DOI:** 10.64898/2026.02.03.703292

**Authors:** Vinay K. Sharma, Nelyda Gonzalez-Almeyda, Simoun Mikhael, Ryan H. Cho, Joseph Aceves, Kristin Ishaya, Sun Woo Kim, Abby Wiesenthal, Rebecca Benhaghnazar, Amirhesam Babajani, Ravinder Abrol, Harry B. Gray, Zeev Gross, Lali Medina Kauwe

## Abstract

Assemblies that combine chemotherapeutics with tumor-targeting proteins are promising agents for treating resistant cancers but require full biochemical characterization before therapeutic deployment. We developed and optimized a HER3-targeting capsomere, HPK2.0, which forms stable nanoscale assemblies with cytotoxic corroles via electrostatic neutralization and shape complementarity. These nanocomplexes exhibit durable serum stability, HER3-dependent tumor invasion, and efficient endosomal escape, resulting in potent and selective cytotoxicity in triple-negative breast cancer (TNBC) cells. In an orthotopic metastatic TNBC model, systemic treatment with HPK2.0–corrole assemblies achieved 67–83% tumor regression, near-complete suppression of spontaneous lung metastasis, and a ∼2-fold improvement in survival relative to mock treatment, with minimal off-target toxicity. By integrating tumor specificity with therapeutic potency, this next-generation protein–corrole platform establishes a clinically scalable strategy for treating metastatic HER3-positive TNBC.

**Significance:** Triple-negative breast cancer (TNBC) is an aggressive disease with high rates of metastasis and mortality, largely because it lacks molecular targets for precision therapy. As a result, patients rely primarily on chemotherapy, which causes systemic toxicity and frequently fails to control metastatic spread. Here, we introduce a targeted therapeutic strategy in which a bioengineered protein selectively recognizes a receptor highly expressed in metastatic TNBC and delivers a potent cytotoxic payload directly into tumor cells. In mouse models, this approach produced robust tumor regression, markedly reduced lung metastases, extended survival, and showed minimal off-target toxicity. These findings establish a versatile platform for targeted treatment of TNBC and highlight a strategy that may be broadly applicable to other HER3-expressing cancers.

## Introduction

Cancer remains a leading cause of death worldwide, and few settings expose the limits of current treatments more starkly than triple-negative breast cancer (TNBC). Defined by the absence of estrogen receptor, progesterone receptor, and HER2 amplification, TNBC has resisted the targeted therapy advances that transformed other breast cancer subtypes. As a result, treatment remains dominated by cytotoxic chemotherapy, where dose limiting systemic toxicity and incomplete tumor control remain persistent barriers to durable benefit (1) These challenges underscore the need for therapeutic strategies capable of achieving tumor-selective lethality while minimizing collateral damage to normal tissues.

One approach to widening the therapeutic window is to decouple cytotoxic potency from indiscriminate exposure by delivering highly active agents selectively to tumor cells and within the intracellular compartments where they exert their effects. Bio-assembled protein delivery systems are well-suited to this goal because their architectures can be programmed to integrate receptor recognition, cellular uptake, and endosomal escape, while accommodating payloads that are too reactive or too membrane impermeant to be safely administered on their own. Building on this concept, we previously developed a heregulin modified cell-targeting protein multimer or capsomere, HerPBK10, now defined as HPK1.0, that noncovalently assembles with sulfonated gallium corroles. This system achieved potent tumor-selective cytotoxicity *in vivo* at doses as low as 8 µg/kg while sparing normal tissues and avoiding cardiotoxicity (2–5). The intense and durable fluorescence of the corrole cargo further enabled direct visualization of tumor accumulation, establishing protein corrole assemblies as intrinsically theranostic agents that unite detection and therapy within a single platform.

Although these initial *in vivo* studies were performed in HER2 amplified tumors, the molecular basis of targeting is ligand engagement of the heregulin receptor axis, with HER3 serving as the primary binding receptor and HER2 acting as a preferred dimerization partner that enhances formation of signaling competent receptor complexes. Thus, what appeared phenotypically as HER2 tumor targeting is more accurately understood as ligand-directed delivery to HER3-enriched receptor assemblies. HER3 ligand–directed engagement by HerPBK10 triggers rapid endocytosis and enables cytoplasmic delivery of associated cargo, including membrane-impermeant macromolecules and sulfonated corroles, in response to endosomal acidification (2, 5–7). This mechanistic understanding substantially broadens the translational relevance of the platform, as HER3 expression extends well beyond HER2 amplified disease.

HER3 is frequently expressed in TNBC and has been implicated in aggressive behavior, metastatic progression, and resistance to therapy, yet it remains difficult to exploit using conventional small-molecule or antibody-based approaches due to its limited intrinsic kinase activity. (8–10) At the same time, translation of first-generation protein corrole assemblies revealed broader design constraints that cannot be addressed through incremental modification alone. Clinical scalability requires efficient and reproducible production of structurally homogeneous capsomeres with preserved biological activity. In parallel, although the relatively large molecular size of corroles (approximately 1000 Daltons) contributes to their stability and cytotoxic potency, it also imposes constraints on pharmacokinetics and bioavailability that must be addressed through coordinated carrier and payload design.

Here, we describe HPK2.0, a next-generation capsomere optimized for HER3-directed tumor targeting, intracellular cargo delivery, and enhanced production efficiency. HPK2.0 incorporates improved production yield, enhanced structural stability, and optimized receptor engagement, and is paired with redesigned corrole cargo to form stable, shape-complementary bio-assemblies. These assemblies selectively invade HER3-expressing triple-negative breast cancer tumors, promote receptor-mediated intracellular trafficking, and enable efficient cytotoxic payload delivery. Together, these advances establish a receptor-defined, modular bio-assembly platform that extends the therapeutic potential of corrole-based agents and provides a generalizable strategy for precision treatment of metastatic cancer.

## Results and Discussion

### HPK2.0 exhibits favorable characteristics for translational development

The bioengineered tumor-invading protein HerPBK10 (HPK1.0) (5) exploits HER3 to penetrate breast tumors resisting ErbB receptor family inhibitors (2). HPK1.0 contains the receptor binding region of the HER3 ligand, neuregulin-1α1 (comprised of Ig-like and EGF-like domains) (11) fused to a membrane-penetrating moiety derived from the adenovirus (Ad) capsid penton base protein modified with a decalysine (K10) tail (**Supplemental Fig. S1**). The penton base drives the formation of pentameric ring or barrel-like structures resembling viral capsomeres (**Supplemental Fig. S1**) with a pH-sensing solvent accessible pore in the center of the pentamer barrel (5). This structure provides pH-mediated endosomolysis after endocytosis (5), enabling access to the cytoplasm. The K10 tail binds anionic cargo whose electrostatic neutralization allows the capsomeres to converge and fit together by shape complementarity, forming a serum-stable polyhedral shell, or nano-capsid, around the cargo (5, 7).

HPK2.0 was developed with a focus on improving protein manufacturability and yield, thereby enhancing clinical translation feasibility. Biological studies of the neuregulin-β isoform show that the EGF-like domain alone is sufficient for receptor binding (12), allowing us to remove the Ig-like region from the targeting ligand and reduce the ligand size by half. The targeting ligand of HPK1.0 extends from the perimeter of the penton base barrel (**Supplemental Fig. S1**), driving the ligands to bend toward the capsomere upon assembly with cargo and potentially restricting their movement on the bio-assembled particle. In the design of HPK2.0, the ligands have been moved to the “top” of the capsomere replacing Arg-Gly-Asp loops in the penton base that are dispensable for receptor binding (**Supplemental Fig. S2**). MD simulation shows that capsomere folding remains unaffected while the ligands enjoy flexible movement (**Supplemental Movie 1**, ligands shown in gold color) and improved solvent exposure on the particle surface compared to HPK1.0, which currently drives the ligands to poke out between capsomeres. MD simulation of HPK1.0 also showed that the K10 may fold into the capsomere pore of HPK1.0 (**Supplemental Figure S3**), thus reducing the potential number of cargo that could be loaded. To alleviate this, we added a spacer between the penton base C-terminus and the K10 tail to sustain solvent exposure of the K10 (**Supplemental Movie 2**, K10 tails shown in red color). Finally, removal of a 50 aa unstable region at the amino [N]-terminus of the penton base yielded a shorter linear sequence (**Supplemental Fig. S2**).

Computational modeling of the HPK2.0 capsomere structure reveals a more compact conformation compared to HPK1.0 (**Fig. 1A**), which may better facilitate packing of the capsomeres around cargo. The smaller capsomere size of HPK2.0 is evident by transmission electron microscopy (TEM), which reveals a considerably smaller ring-shaped structure compared to HPK1.0 (**Fig. 1B**), consistent with the computationally predicted model. Denaturing gel electrophoresis of HPK2.0 shows an improved purity compared to HPK1.0 (**Fig. 1C**), which is reflected by the improved polydispersity index of 2.0 compared to 1.0 (**Fig. 1D**). Electrophoretic mobility shift assay (EMSA) shows that HPK2.0 better immobilizes nucleic acid cargo compared to HPK1.0 (**Fig. 1E**). When each is assembled with a fluorescently tagged oligonucleotide cargo to form nano-bioparticles (NBPs), 2.0 facilitates augmented binding to HER3 positive cells with greater receptor specificity (confirmed by excess competing ligand neuregulin or NRG) compared to 1.0 (**Fig. 1F**). When delivering the chemotherapy drug doxorubicin (Dox) by intercalation in oligonucleotide duplex cargo, 2.0 showed *three orders of magnitude improved IC50* (36 nM) compared to 1.0 (3.8 µM; **Fig. 1G**). Taken together, the modified protein sequence not only improved yield and purity, but also enhanced cargo loading, receptor specificity, and selective potency on HER3-overexpressing tumor cells.

**Figure 1.**
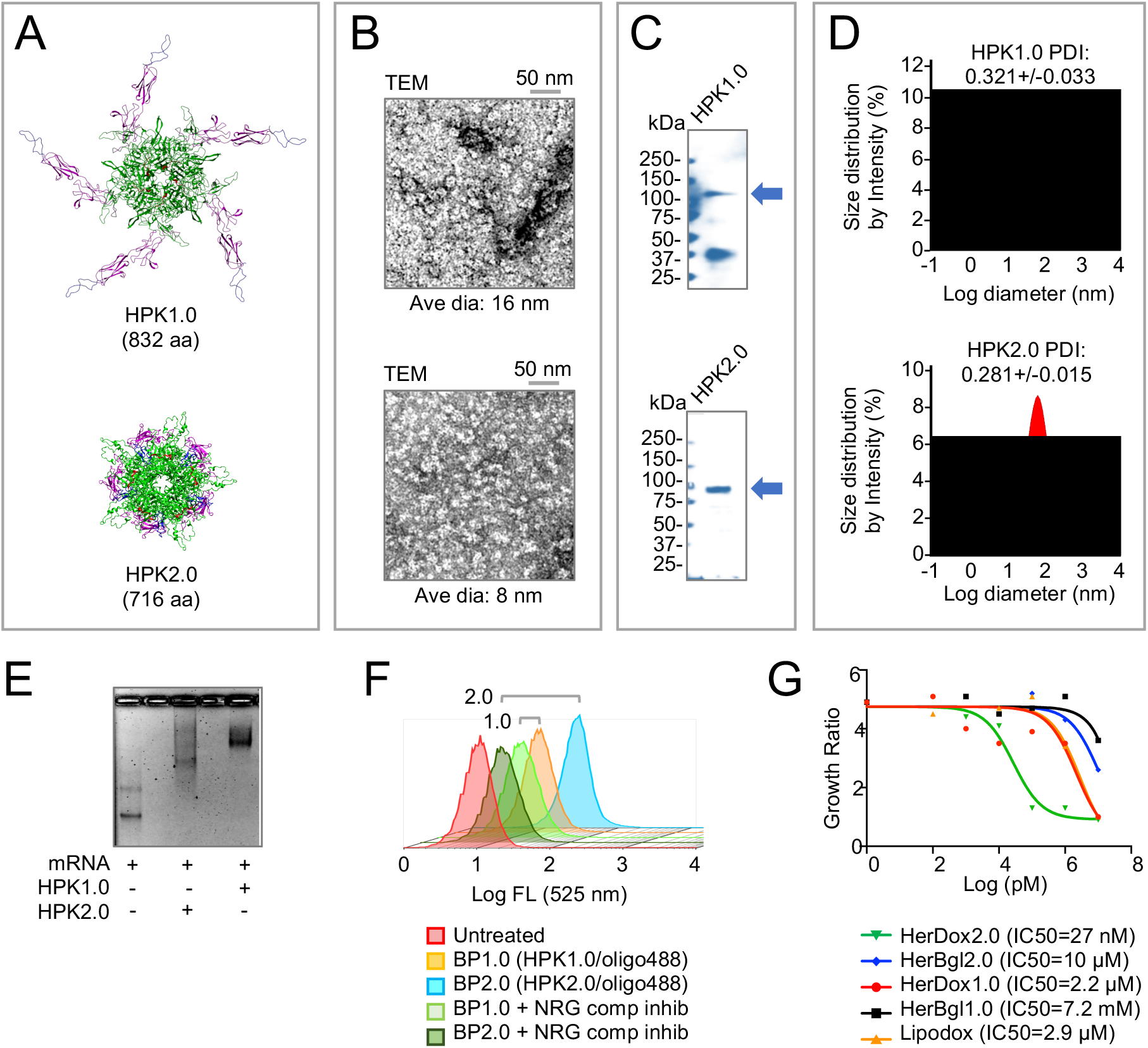
HPK1.0 and HPK2.0 protein comparisons. **A**, Computational modelling of capsomeres. Functional domains are delineated by color: HER3 targeting ligand (magenta), penton base (green), and cargo loading (red). **B**, Transmission electron microscopy (TEM) of capsomeres. **C**, SDS-PAGE of denatured protein after affinity purification. Arrow delineated full-length protein. **D**, Dynamic light scattering of capsomeres. **E**, Electrophoretic mobility shift analysis (EMSA) of nucleic acid cargo (1 kb mRNA, 0.5 µg) -/+ each corresponding protein (pre-incubated at 0.01 mRNA:protein molar ratio, 30 min, room temp before gel electrophoresis). **F**, Fluorescence count of 4T1 mouse TNBC cells -/+ incubation with indicated fluorescently tagged bioparticles -/+ competing ligand (NRG). **G**, Cytotoxicity curves of 4T1 cells exposed to titrating concentrations of indicated bioparticles delivering a tumoricidal drug (Dox) or inert cargo (Bgl), compared to liposomal doxorubicin (Lipodox).

### Phosphorus corrole properties favor therapeutic translation

Ideally, small-molecule drugs typically have molecular weights of less than 500 Da and a simple chemistry with few functional groups (13–15). Here we examined whether the theranostic benefits of S2Ga [Mw = 1021 Da], including potent tumoricidal activity combined with fluorescence emission (7, 16), can be sustained while reducing the chemical size and complexity. Accordingly, we explored gallium and phosphorus complexes of the most recently introduced *meso*-C-carboxylated corrole (14) and examined the effect of the macrocycle by comparing S2Ga with (tcc)Ga [Mw = 494 Da]; and examined the effect of the central metal by comparing (tcc)Ga with (tcc)P(OH)_2_ [Mw = 489 Da] (**Fig. 2**). Whereas all three compounds exhibited water solubility, (tcc)P(OH)_2_ showed the greatest degree of aqueous distribution (logP = -2.35; **Table 1**). In phosphate-buffered saline (PBS), (tcc)P(OH)_2_ exhibited blue-shifted absorption and a threefold higher extinction coefficient compared to S2Ga or (tcc)Ga (**Table 1 and Supplemental Fig. S4**). Employing fluorescein as a reference (17), we found that (tcc)P(OH)_2_ emitted a higher fluorescence quantum yield (12%) compared with S2Ga (8%) and (tcc)Ga (3%). Taken together, (tcc)P(OH)_2_ exhibits improved physicochemical characteristics compared to the other two corroles.

**Figure 2.**
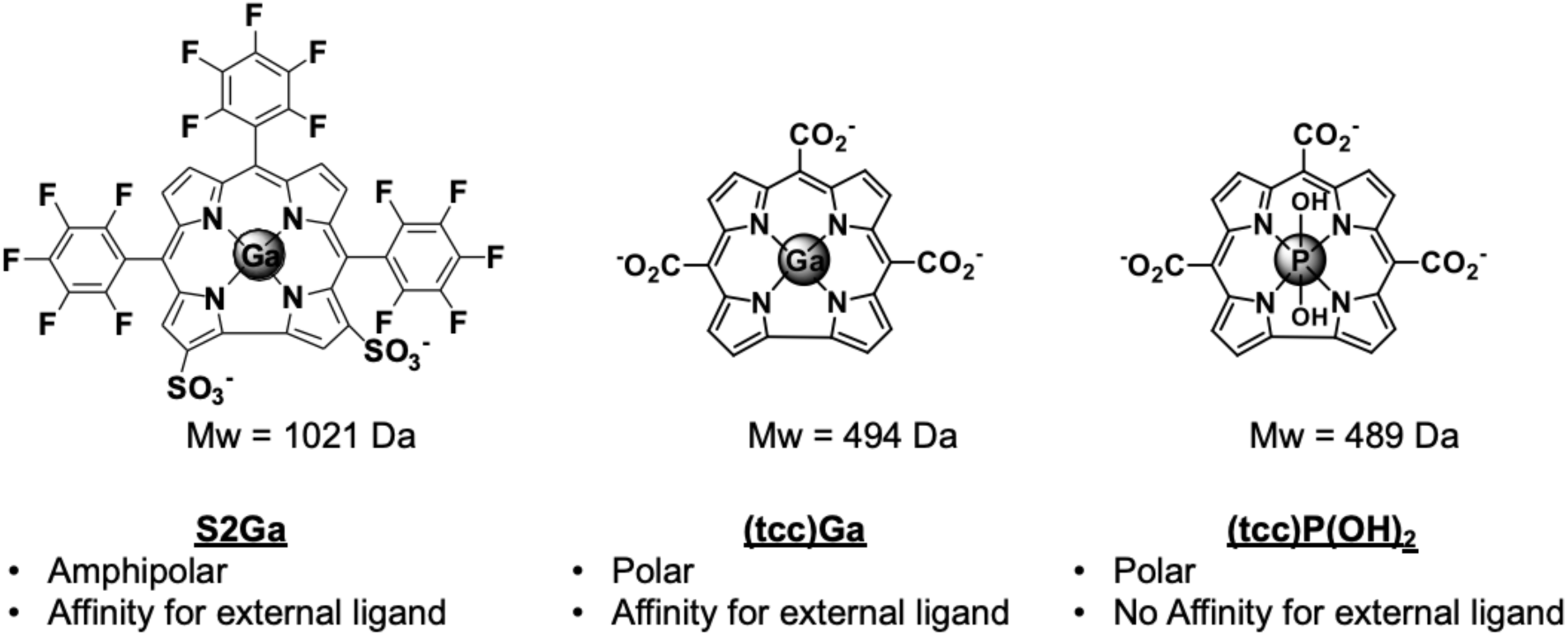
Structures of corrole compounds used in this study. Chemical structures of the previous lead molecule S2Ga and the recently developed corroles (tcc)Ga and (tcc)P(OH)₂ with key features are shown.

**Table 1.** Summary of the photophysical and other relevant properties* of the corroles.

| Compounds | Absorbance<br>($\lambda_{\text{max}}$) in nm | Emission<br>($\lambda_{\text{max}}$) in nm | Extinction<br>Coefficient<br>($\epsilon$) in $\text{M}^{-1} \text{cm}^{-1}$ | Quantum<br>Yield<br>($\phi_F$) | Partition<br>Coefficient<br>(logP) |
| --- | --- | --- | --- | --- | --- |
| S2Ga | 424 | 627 | 74,700 | 0.08 | 0.72 |
| (tcc)Ga | 417 | 612 | 78,800 | 0.03 | -1.95 |
| (tcc)P(OH)<sub>2</sub> | 401 | 587 | 216,600 | 0.12 | -2.31 |
| Fluorescein | 492 | 517 | ** | 0.75 | ** |

\* All measurements were conducted in 10 mM PBS at pH 7.4.
\*\* Data not available for some entries.

Isothermal titration calorimetry (ITC) was used to evaluate the binding capacity of each corrole to HPK2.0. All three corroles exhibited thermodynamically favorable binding to HPK2.0 with the reaction being endothermic for S2Ga and exothermic for the other two corroles (**Fig. 3, A–B** and **Supplemental Fig. S5**). The minimally effective concentrations revealed by these binding curves indicate that maximal assembly occurs at a corrole:HPK2.0 ratio of 20-50 molecules of corrole bound per HPK2.0 monomer (**Table 2**), in agreement with our previous studies in which we have measured corrole content in particle preparations based on absorbance and fluorescence measurements (2, 3, 18). Accordingly, we used this stoichiometry to prepare bio-assemblies (see *Materials and Methods* for experimental details).

**Figure 3.**
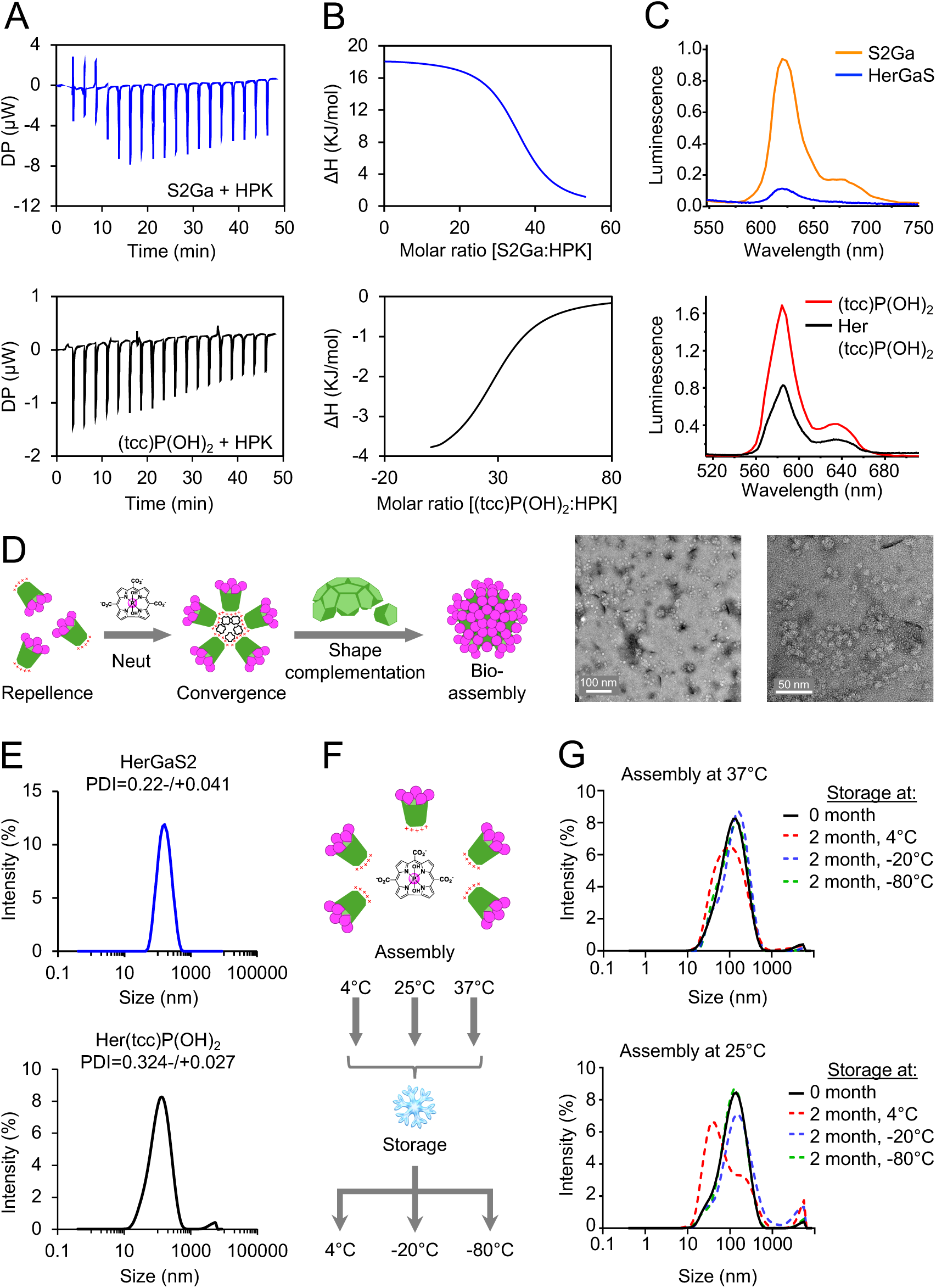
Characterization of HPK2.0–corrole bio-assemblies. **A–B**, Isothermal titration calorimetry (ITC) analyses (showing thermograms and binding curves) of HPK2.0 reacting with S2Ga and (tcc)P(OH)_2_. **C**, Photoluminescence spectra of unbound (black traces) and HPK2.0-bound (red traces) S2Ga, (tcc)Ga, and (tcc)P(OH)₂. **D**, Schematic of HPK2.0 self-assembly with corroles, showing charge-repellence of capsomeres in solution that is counteracted by the addition of anionic corroles that neutralize the charges and allow capsomeres to converge by shape complementation, forming polyhedral bio-assemblies. Insets show transmission electron microscopy (TEM) of Her(tcc)P(OH)_2_ particles at two different magnifications. **E**, Dynamic light scattering (DLS) analyses showing size distributions of the bio-assemblies. **F**, Schematic showing differential assembly and storage conditions used for evaluating particle stability. **G**, Hydrodynamic diameters of Her(tcc)P(OH)_2_ particles assembled and stored at indicated temperatures.

**Table 2.** Thermodynamic properties of protein-corrole bioassemblies.

| Bioassembly name | $K_d$ (μM) | $\Delta H$ (kJ/mol) | $\Delta G$ (kJ/mol) | $-T\Delta S$ (kJ/mol) | N (sites) |
| --- | --- | --- | --- | --- | --- |
| HerGaS2 | 2.3 | +18.4 | -32.2 | -50.6 | 35 |
| Her(tcc)P(OH)<sub>2</sub> | 13.5 | -4.3 | -27.8 | -23.6 | 30 |

Protein binding resulted in the least photoluminescence quenching for tccP(OH)₂, relative to the other corroles (**Fig. 3C**; **Supplemental Fig. S6**). Notably, HPK2.0 retained S2Ga and tccGa more strongly (**Supplemental Fig. S7**), suggesting that the absence of a central metal in tccP(OH)₂ limits its interaction with the protein to primarily electrostatic and van der Waals forces. Under these conditions, the corroles are likely bound predominantly to the K10 tail, promoting convergence of HPK protomers and encapsulation of the corroles within the nano-capsid (**Fig. 3D**). This model is supported by TEM imaging of bio-assemblies formed from mixing HPK2.0 with (tcc)P(OH)_2_ which reveal well-defined polyhedral particles with an average diameter of ∼20 nm (**Fig. 3D**). By contrast, the poor colloidal stability of tccGa, evidenced by precipitation upon mixing with HPK2.0 (**Supplemental Fig. S7**), precluded its further evaluation.

The remaining bio-assemblies, designated HerGaS2 and Her(tcc)P(OH)_2_, formed hydrodynamic diameters ranging between 20-40 nm (**Fig. 3E**). Particle stability was examined by forming the assemblies at 4 °C, 25 °C, or 37 °C, followed by storage under extended times (2 months, 1 year) and temperatures (4 °C, –20 °C, –80 °C) (**Fig. 3F**). Stability was assessed by looking at any time and temperature-dependent aggregation that would be indicated by an increase in hydrodynamic diameter. Assemblies prepared at 37 °C remained stable for up to 2 months under all storage conditions (4 °C, –20 °C, –80 °C) (**Fig. 3G**) and up to 1 year under storage at -20°C and -80°C but not 4 °C (**Supplemental Fig. S8**). In contrast, assemblies generated at 25 °C were stable only at –20 °C or –80 °C (**Fig. 3G**) and precipitated after storage at 4 °C, whereas those formed at 4 °C aggregated under all conditions (data not shown). Together, these findings demonstrate that assembly at 37 °C produces optimal Her(tcc)P(OH)_2_ formulations with excellent long-term stability.

### Her(tcc)P(OH)₂ assemblies reduce tumor growth and metastasis in HER3^+^ TNBC

To investigate the tumoricidal potential of the protein–corrole bio-assemblies, growth inhibition assays were performed using 4T1 mouse TNBC cells and were compared to activities on NIH/3T3 mouse fibroblast cells. Among the tested bio-assemblies, Her(tcc)P(OH)₂ exhibited the strongest cytotoxicity against 4T1 cells (IC_50_ = 7.56 µM; **Supplemental Table 1**), resulting in significantly reduced cell viability by 24 hours **(Fig. 4A–B**). Its weaker impact on NIH/3T3 cells (IC_50_ = 53 mM; **Supplemental Table 2**) suggests that Her(tcc)P(OH)₂ has preferential selectivity for the cancer cells. Although HER3 is detectable in lysates from both cell lines, surface expression differs markedly: 4T1 cells display HER3 at levels comparable to human TNBC (MDA-MB-468), melanoma (MDA-MB-435), and HER2⁺ breast cancer cells (BT-474L), whereas NIH/3T3 cells express an order of magnitude less (**Supplemental Fig. S9**). Hence, tumor preference is likely mediated by the tropism of HPK2.0 for HER3. This is supported by the absence of any notable effect of (tcc)P(OH)₂ alone on both cell types (**Fig. 4, C–D**), indicating that delivery by HPK2.0 is crucial for therapeutic activity. The requirement for HPK2.0-mediated delivery is further supported by the activities of HerGaS2 bio-assemblies which also showed preferential cytotoxicity on 4T1 cells compared to NIH-3T3 cells (**Fig. 4B**) and contrasted with S2Ga alone, which showed negligible to absent activity on either cell type (**Fig. 4D**). Importantly, empty particles (HerLLAA2.0) lacked any notable effect on cell density (**Fig. 4B**), indicating that the therapeutic effect is primarily due to the corrole cargo. Of further note is that, both Her(tcc)P(OH)₂ and HerGaS2 assemblies displayed augmented cytotoxicity upon photoexcitation, consistent with the intrinsic photodynamic properties of corroles (**Supplemental Fig. S9 and Table 1–2**) (19–22).

**Figure 4.**
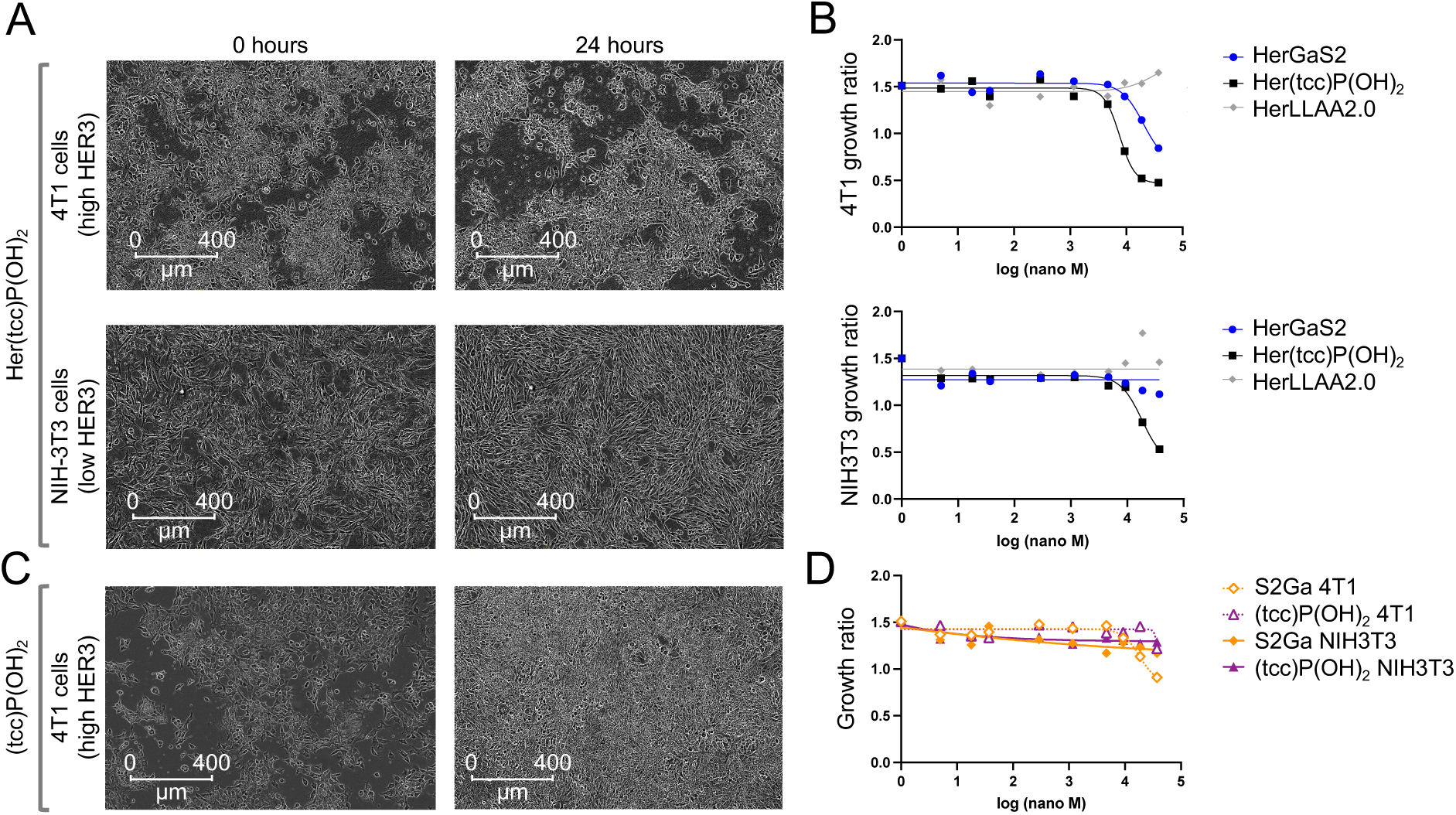
Cell morphology and growth response to HER3-targeted corrole conjugates. **A**, Brightfield images of 4T1 (high HER3) and NIH-3T3 (low HER3) cells at 0 and 24 h following treatment with Her(tcc)P(OH)_2_. Scale bars, 400 µm. **B**, Growth ratio of 4T1 (top) and NIH-3T3 (bottom) cells as a function of corrole concentration for HerGaS2, Her(tcc)P(OH)_2_, and HerLLAA2.0. **C**, Brightfield images of 4T1 cells at 0 and 24 h following treatment with (tcc)P(OH)_2_. Scale bars, 400 µm. **D**, Growth ratio of 4T1 and NIH-3T3 cells as a function of corrole concentration for S2Ga and (tcc)P(OH)_2_.

Given the *in vitro* structural stability of Her(tcc)P(OH)₂ assemblies and its tumor cell preferential cytotoxicity, we next examined whether these advantages would extend to *in vivo* tumor targeting and therapeutic efficacy. Specifically, we sought to determine whether the HER3-directed tropism of HPK2.0 could mediate selective tumor localization and antitumor activity in immunocompetent (BALB/c) mice bearing orthotopic HER3^+^ 4T1-Luc tumors, an aggressive and spontaneously metastatic model of TNBC.

To evaluate tumor tropism *in vivo*, we examined the time-sequential tissue distribution following a single intravenous (tail vein) administration of Her(tcc)P(OH)_2_ in both tumor-bearing and non–tumor-bearing mice. Her(tcc)P(OH)_2_ exhibited pronounced tumor accumulation with the tumor content of corroles (150-210 nmoles/g tissue) increasing from 2 to 24 hours post-injection (**Fig. 5A**). In contrast, uptake in clearance organs such as the liver and spleen was present but remained significantly lower (<90 nmoles/g tissue) relative to the corrole content in tumors at all time points measured. In non–tumor-bearing controls, Her(tcc)P(OH)_2_ distributed primarily to the liver, spleen, and kidneys, consistent with normal clearance pathways but still remained well below 90 nmoles corrole/g tissue (**Fig. 5B**). Taken together, these findings demonstrate that Her(tcc)P(OH)_2_ predominantly localizes to tumors *in vivo*, while non-tumor uptake is significantly lower, in both tumor and non-tumor bearing animals.

**Figure 5.**
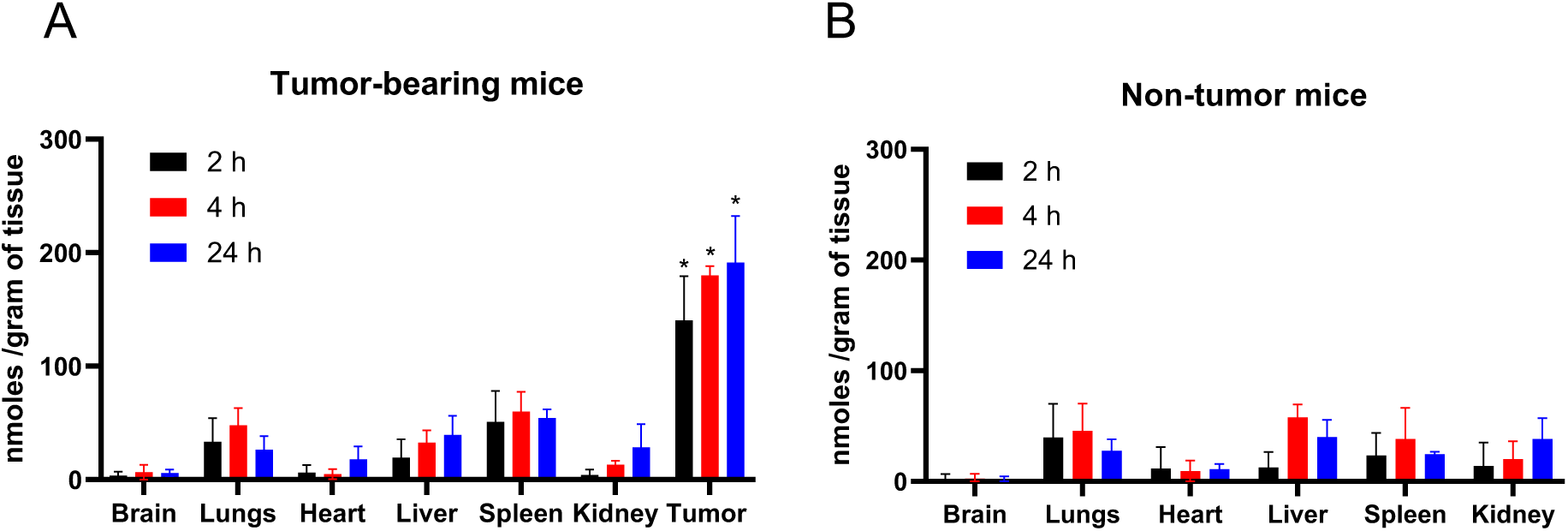
Time-sequential tissue distribution of HPK2.0–corrole nano-assemblies. Graphs show tissue content of (tcc)P(OH)_2_ in (**A**) tumor-bearing and (**B**) non–tumor-bearing mice following a single intravenous injection of Her(tcc)P(OH)₂ equating 12.5 nmol (tcc)P(OH)_2_. Error bars represent mean ± SD (n=3 per time point). * p=the following indicated values comparing tumors with corresponding non-tumor organs at each time point (Dunnett’s multiple comparisons test): At 2 h, tumor uptake was significantly higher than in lungs (*p* = 0.0251), liver (*p* = 0.0330), and spleen (*p* = 0.0162); At 4 h, tumor uptake was significantly higher than in the brain (*p* = 0.0013), lungs (*p* = 0.0265), heart (*p* = 0.0245), liver (*p* = 0.0288), spleen (*p* = 0.0326), and kidney (*p* = 0.0008); At 24 h, tumor uptake was significantly higher than in the brain (*p* = 0.0385), lungs (*p* = 0.0403), heart (*p* = 0.0304), and kidney (*p* = 0.0383); differences vs liver (*p* = 0.1157) and spleen (*p* = 0.0885) were not significant.

We next evaluated *therapeutic efficacy* in mice bearing established orthotopic TNBC tumors, following an administration regimen based on the minimally effective concentrations determined in cell-based assays combined with the time-sequential tissue distributions previously determined *in vivo* (described in Methods; **Fig. 6A**) (7). We applied Response Evaluation Criteria In Solid Tumors (RECIST) adapted for mouse subjects (23) to evaluate primary tumor response based on four categories: progressive disease (>20% increase in tumor size); stable disease (<20% increase in tumor size and <30% decrease in size); partial response (at least 30% decrease in tumor size); and complete response (disappearance of all lesions). Based on these criteria, mice receiving systemic delivery of Her(tcc)P(OH)_2_ showed the highest regression of primary tumors among all cohorts with nearly 30% of the cohort showing complete responses, only one mouse with progressive disease, and the rest displaying partial response (**Fig. 6B–C**). HerGaS2 treatment also reduced tumor growth eliciting partial response in 67% of mice although the remaining 33% showed progressive disease and none with complete response (**Fig. 6B–C**). The drug-free control, HerLLAA2.0, elicited no marked differences from mock (saline) treatments (**Fig. 6B–C)**, suggesting that the tumoricidal activity of the bio-assemblies is elicited by the corrole cargo. In partial agreement with this, free corroles, (tcc)P(OH)_2_ and GaS2, elicited tumor regression or suppression in 70% of each cohort but nearly 30% in each still showed progressive disease (**Supplemental Fig. S10**).

**Figure 6.**
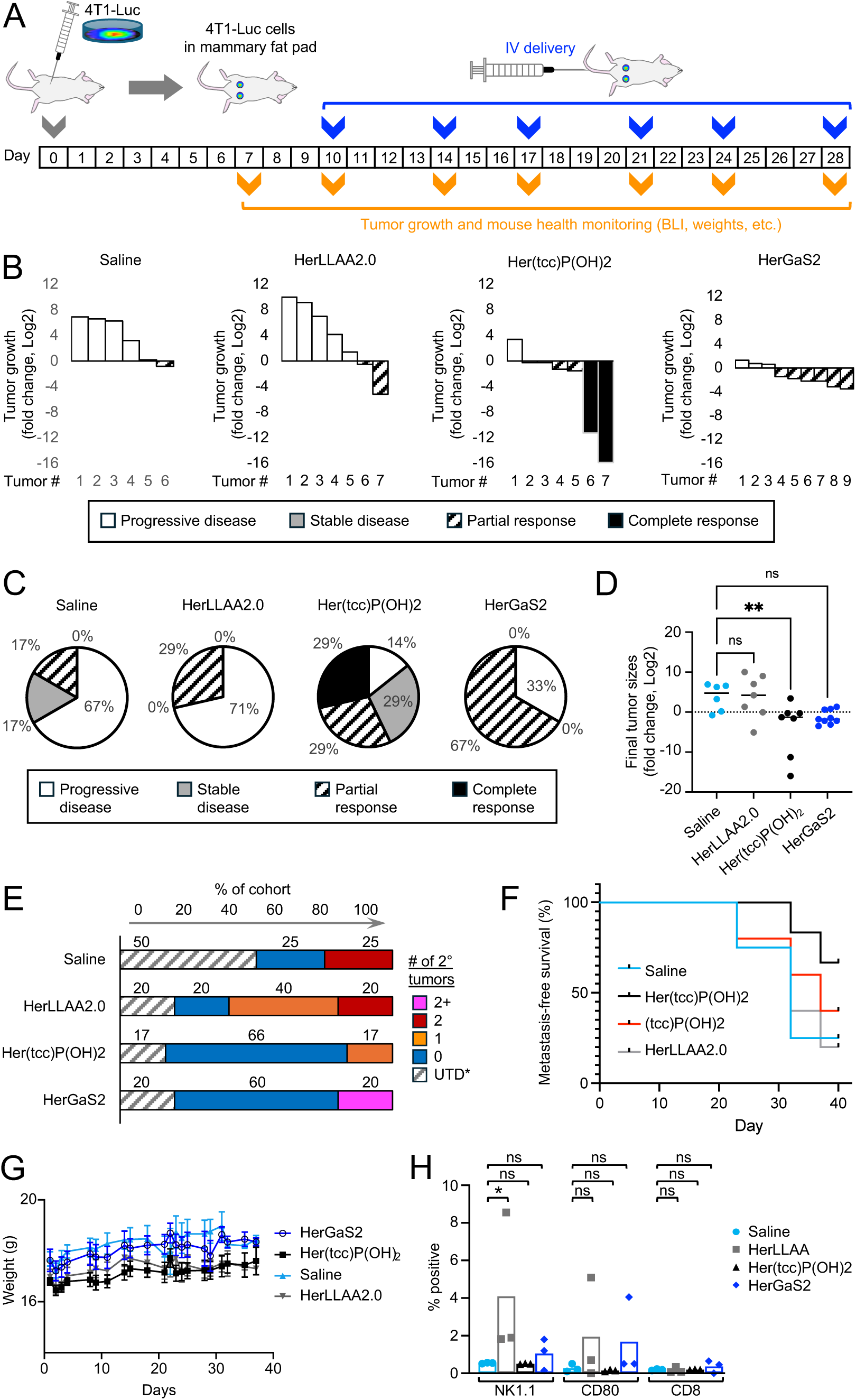
Therapeutic response, metastatic outcomes, and tolerability of HER3-targeted corrole bio-assemblies in an orthotopic TNBC model. **A,** Experimental timeline illustrating orthotopic implantation of 4T1-luc cells into the mammary fat pad of BALB/c mice and subsequent intravenous (i.v.) administration of saline or the bio-assemblies. Tumor growth and treatment responses were monitored over time. **B,** Waterfall plots showing log_2_ fold change in primary tumor growth for individual tumors following treatment with HerLLAA2.0, Her(tcc)P(OH)_2_, or HerGaS2 at a dose of 0.22 mg/kg compared with saline controls. Tumors are classified according to RECIST-adapted response criteria: progressive disease, stable disease, partial response, or complete response. **C,** Pie charts summarizing the distribution of RECIST-adapted therapeutic response categories for each treatment cohort. Percentages indicate the fraction of tumors in each category. **D,** Scatter plot of log_2_ fold change in final primary tumor sizes for individual tumors across treatment groups. Horizontal solid lines indicate group means. Statistical comparisons are shown as indicated (ns, not significant; **p < 0.01). **E,** Distribution of the number of secondary tumors per mouse across treatment cohorts. *UTD, unable to determine (i.e. due to early death or required euthanasia early in the experiment). **F,** Kaplan–Meier analysis of metastasis-free survival for mice treated with saline, HerLLAA2.0, HerGaS2, or Her(tcc)P(OH)_2_. **G,** Body weight measurements over time for mice receiving the indicated treatments. Data are shown as mean ± SEM. **H,** Quantification of tumor-associated immune cell markers (NK1.1^+^, CD80^+^, and CD8^+^) across treatment groups. Each symbol represents an individual mouse; bars indicate mean ± SEM. *Cohort sizes ranged from n = 4–6 mice per treatment group*.

Her(tcc)P(OH)_2_ treatment not only elicited the highest overall regression in primary tumor sizes compared to the other treatments (**Fig. 6D** and **Supplemental Fig. S10**) but also the lowest number of secondary tumors among the cohorts (**Fig. 6E**), which include metastases detected in the lungs, peritoneum, and other sites. This comes into play by the Her(tcc)P(OH)_2_-treated group presenting the highest metastasis-free survival among the cohorts (**Fig. 6F** and **Supplemental Fig. S11**). All mice appeared to tolerate the treatments and showed no signs of significant weight loss that would otherwise indicate adverse effects or toxicity compared to mock (saline) treated controls (**Fig. 6G** and **Supplemental Fig. S12**). Evaluation of immune-activation markers (NK1.1, CD80, and CD8^+^) delineating natural killer cells, CD80+ macrophages, and CD8+ T cells showed no significant differences among Her(tcc)P(OH)_2_, HerGaS2, and saline groups (**Fig. 6H** and **Supplemental Fig. S13**), suggesting that the assemblies do not recruit an immune-mediated response.

## Conclusion

TNBC presents a persistent therapeutic challenge due to its molecular heterogeneity, aggressive metastatic behavior, and lack of druggable receptors. Here, we report HPK2.0–corrole bio-assemblies as a next-generation HER3-targeting platform that integrates rational protein engineering, receptor recognition, shape-complementary nanoparticle assembly, and endosomal escape into a unified therapeutic architecture. These assemblies exhibit durable cargo encapsulation, selective tumor penetration, and potent cytotoxicity, achieving 67–83% regression of established primary tumors and near-complete suppression of spontaneous lung metastases in an orthotopic TNBC model, translating to approximately a two-fold improvement in survival relative to controls while sparing normal tissues. Beyond corrole delivery, HPK2.0 demonstrates platform generalizability: when delivering doxorubicin via oligonucleotide cargo, the capsomere increased drug potency by three orders of magnitude compared to the first-generation HPK1.0, highlighting that efficacy is governed by capsomere architecture and HER3 engagement rather than payload chemistry alone. By linking molecular design principles to functional outcomes *in vivo*, these studies establish HPK2.0 as a modular, mechanistically informed nanotherapeutic platform capable of overcoming heterogeneity and metastatic progression in HER3-expressing cancers. This work provides a foundation for further translational development of protein-guided, intracellular delivery systems for TNBC and other receptor-defined malignancies.

## Materials and Methods

### Construction of HPK2.0

The HPK1.0 DNA construct is comprised of the pRSET-A plasmid vector containing the coding sequence of HerPBK10(1, 3) cloned in-frame to produce a recombinant fusion protein comprising the amino acid sequence shown in **Supplemental Fig. S1**. The HPK2.0 DNA construct (**Supplemental Fig. S2**) differs from HPK1.0 by: removing the Ig-like region from the targeting ligand, relocating the targeting ligand to replace a solvent exposed flexible Arg-Gly-Asp loop in the penton base domain and flanking the ligand by neutral (Gly-Gly-Ser)*2 residues serving as flexible linkers; extending the K10 tail by the insertion of a (Gly-Gly-Ser) flexible linker to enhance exposure and interaction with cargo; and truncating the penton base domain by removing a 50 amino acid unstable region at the amino [N]-terminus of the penton base.

### Implicit Molecular Dynamics Simulation of Pentameric HPK2.0

The initial protein structure was obtained from a previously prepared PDB file generated in house. Protonation states of all ionizable residues were assigned at pH 7.0 using the PDB2PQR web server (24). The pentameric HPK2.0 was relaxed to assess its stability and structural dynamics using the pmemd.cuda (GPU) program from the AMBER simulation suite (25) with the ff14SBonlysc force (26), employing a generalized Born implicit solvent model for all stages of the simulation. Step 1, the system was energy-minimized with backbone restraints to relax unfavorable contains; step 2, the protein was heated from 10K to 310.15 K over 0.5 ns to enable the system to adjust to physiological temperature; step 3; the unrestrained system was equilibrated for 1 ns to allow protein to fully relax within the implicit solvent environment; step 4, the equilibrated system was simulated for 100 ns to generate the production trajectory used for analysis.

### Transmission electron microscopy (TEM)

Homogenous protein samples were prepared for transmission electron microscopy (TEM) by dispersion in hexane to achieve a uniform sample onto a carbon coated grid. The fixed samples of protein HPK 1.0 and HPK 2.0 were imaged using a TF20 (FEI Tecnai) transmitting electron microscope (Electron Imaging Center for NanoMachines, California NanoSystems Institute, UCLA). Images were acquired using a TIETZ F415MP CCD camera and processed using TIETZ Tomography package.

### Dynamic Light Scattering (DLS)

Dynamic Light Scattering (DLS) was performed to determine the average particle size of HPK 1.0 and HPK 2.0 proteins using a Malvern Zetasizer. A sample of 20 µg of protein was loaded into a quartz cuvette. The size analysis consisted of 7 measurements per sample, each measurement consisting of 45 runs. The 7 measurements were averaged to yield the most frequent particle size while accounting for intensity fluctuations in the sample.

### Electrophoretic Mobility Shift Assay (EMSA)

Each nucleic acid species (400 ng mRNA encoding GFP; MR700A-1, System Biosciences) was incubated with either HPK1.0 or HPK2.0 at a 100:1 molar ratio for 30 min at room temperature in 1X annealing buffer (10 mM Tris pH 7.5, 50 mM NaCl, and 1 mM EDTA) followed by electrophoresis on a 1.5% agarose gel and post-electrophoresis staining with ethidium bromide following standard procedures.

### Cell binding assay

Indicated adherent cells were grown to 70-80% confluency before detachment and resuspension in Buffer A (0.5 M HEPES (pH7.4), 1 M MgCl2, BSA-final 3%, DMEM with no FBS) at 2,500,000 cells/mL, then distributed into 0.2 mL aliquots (500,000 cells/aliquot) in microfuge tubes that received 0.6 mL volumes of binding mixtures. Binding mixtures included 1XPBS containing HPK1.0 or 2.0 pre-assembled with fluorophore (Alexa488)-tagged oligonucleotide (conc: 1.7 µg/uL (17µM), 0.7 µg/µL (8.75µM) -/+ neuregulin (NRG; R&D systems) at 10x molar excess (compared to HPK concentration). Mixtures were incubated on ice for 1 hour with rocking, then gently pelleted at (200 x G) for 5 minutes at 4 deg C and washed 3 times in ice cold PBS, resuspended in ice cold PBS, and processed by flow cytometry to measure fluorescent events.

### UV-vis spectroscopy

Electronic absorption spectra of the synthesized corroles were recorded on a Cary 8454 UV–vis spectrophotometer (Agilent Technologies) using quartz cuvettes with a 1.0 cm path length. All measurements were conducted in 50 mM PBS (pH 7.4). Extinction coefficients (ε) were determined by plotting absorbance versus concentration and applying the Beer–Lambert law (A = εcl).

### Fluorescence spectroscopy

Steady-state emission spectra of corrole–protein assemblies and free corroles were measured on a fluorometer (FSS, Edinburgh Instruments) equipped with a Xenon arc lamp at 298 K under aerobic conditions. Measurements were performed in 10 mM PBS (pH 7.4) using 1.0 cm quartz cuvettes. Fluorescein in 10 mM PBS (Φ = 0.75) (17) was used as a standard for determining relative quantum yields (Φ). Absorption spectra of both fluorescein and corroles were recorded, and excitation wavelengths were selected at the intersection of their absorption spectra. The applied excitation wavelengths were 424 nm for (tcc)P(OH)₂, 465 nm for (tcc)Ga, and 445 nm for S2Ga.

Quantum yields were calculated using the comparative method according to the following simplified equation:

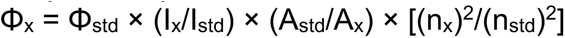

where *I* is the integrated emission intensity, *A* is the absorbance at the excitation wavelength (since excited at the intersection point, A_x_ = A_std_), *n* is the refractive index of the solvent (PBS in both cases, thus n_x_ = n_std_), and subscripts *x* and *std* refer to the sample and fluorescein standard, respectively.

### Partition coefficient (log P) determination

The lipophilicity of the synthesized corroles was evaluated by determining their octanol–PBS partition coefficients (log P) using the shake-flask method. Briefly, known concentrations of the compounds were equilibrated between n-octanol and water phases by vigorous mixing followed by phase separation. The concentration of corroles in each phase was quantified using UV–vis spectroscopy on an Agilent Technologies Cary 8454 spectrophotometer. Quartz cuvettes of 1.0 cm path length were used for all measurements. The partition coefficient was calculated as the ratio of absorbance-derived concentrations in the n-octanol and aqueous phases, expressed as log P.

### Particle assembly

Binding ratios of protein and cargo were determined using a Malvern Panalytical MicroCal PEAQ Isothermal Calorimeter (ITC). HPK protein (3.5 μM) in a stirred cell received cargo in 2 µL titrations at 37°C until binding reached saturation as measured by heat exchange. Analysis was performed using the PEAQ ITC analysis software to determine enthalpy (ΔH), entropic free energy (-TΔS), Gibbs free energy (ΔG), and the dissociation constant (KD) of the system. The protein:cargo stoichiometry identified at the minimally effective concentration (MEC) provided the parameters for particle assembly.

To assemble Her(tcc)P(OH)_2_ particles, (tcc)P(OH)_2_ corrole was slowly added with stirring into an HPK 2.0 solution at the MEC stoichiometries determined by ITC. The mixture of HPK 2.0 and (tcc)P(OH)_2_ (1:30 molar ratio) was incubated at 4°C overnight in NaP (pH 8.0) to produce Her(tcc)P(OH)_2_ complex. Following overnight incubation, the complex underwent ultrafiltration using a 50 K molecular weight cut-off (mwco) membrane to isolate the assembled complex from any unbound components. Retentates were profiled by DLS and quantified based on protein content by measuring optical density (OD) at 280 nm and corrole content by measuring the OD at 418 nm using a Spectramax M2 plate reader (Molecular Devices, LLC).

To assess particle stability, a variety of different assembly and storage conditions were tested. Particle complexes were prepared as described earlier, comparing three different assembly conditions (overnight incubation at 4°C, room temperature for 1 hour, or 37°C for one hour), followed by ultrafiltration and DLS measurements. To assess storage stability, 20 µg aliquots (based on protein content) of each complex were stored at three different temperatures (4°C, -20°C, and –80°C) with DLS profiling performed biweekly at room temperature and 37°C.

### Cell Viability and Cytotoxicity Assay

4T1 triple-negative breast cancer (TNBC) cells and NIH-3T3 fibroblast cells were used to assess the cytotoxicity of protein–corrole bio-assemblies. Cells were seeded in 96-well plates at 1 × 10⁴ cells per well and allowed to adhere overnight in complete growth medium at 37 °C with 5% CO_2_. The next day, cells were treated with protein–corrole assemblies [Her(tcc)P(OH)_2_ or HerGaS2], free corroles [(tcc)P(OH)_2_ or S2Ga], or protein-only controls (HerLLAA2.0) at 10 µM. For dose–response studies, serial dilutions (0.001–100 µM) were prepared in serum-free medium. Treatments were applied for 3 h under controlled light conditions: plates were either covered with foil for dark treatment or exposed to low-intensity white light for 1 h. Medium was then replaced with complete growth medium for recovery. Brightfield images were acquired at 0, 12, and 24 h using the Incucyte® Live-Cell Analysis System (Sartorius) at 37 °C and 5% CO₂. Cell confluence and proliferation were quantified with the Incucyte software and normalized to untreated controls. IC_50_ values were determined by fitting dose–response curves with a four-parameter logistic model in GraphPad Prism.

### HER3 immunodetection

To assess HER3 levels in cell lysates, 4T1 breast cancer and NIH-3T3 fibroblast cells were cultured to ∼80% confluence, detached, and lysed in Triton X-100 containing protease inhibitors. Total protein was quantified using the Pierce™ BCA assay. Equal protein amounts were subjected to SDS–PAGE and western blotting to examine ErbB3 expression. Samples were mixed with 4X Laemmli buffer, denatured at 95 °C for 7 min, briefly centrifuged, and kept on ice before loading. Precision Plus Protein Unstained and All Blue standards (1:1) served as molecular weight markers. Proteins were resolved on precast gels using a Mini-PROTEAN system in 1X SDS running buffer and imaged under stain-free settings on a ChemiDoc MP system prior to transfer. Transfers to nitrocellulose membranes were performed using wet transfer at 25 V for 30 min in transfer buffer containing 20% ethanol. Membranes were blocked for 1 h in either 3% BSA or 5% milk, then incubated overnight at 4 °C with polyclonal anti-HER3/ErbB3 (Cat. No. PA5-14636, Thermo Fisher Scientific, Waltham, MA, USA) at 1:1,000 dilution. After washing with TBS-T and TBS, membranes were incubated with HRP-conjugated polyclonal secondary antibodies (Goat Anti-Rabbit IgG H&L, Cat. No. ab6721; Goat Anti-Mouse IgG H&L, Cat. No. ab6789, Abcam, Cambridge, UK) (1:5,000) for 1 h, washed again, and developed using Pierce ECL substrate. Chemiluminescence was captured using the ChemiDoc MP under high-sensitivity settings.

To assess cell-surface HER3, cells were grown in T75 flasks to 70–80% confluence, washed twice with PBS, detached with Trypsin–EDTA, neutralized in complete medium, pelleted, and resuspended in PBS before viability was confirmed by Trypan Blue exclusion. Approximately 5 × 10⁵ viable cells were transferred into 500 µL ice-cold PBS, pelleted at 200 × g for 5 min at 4 °C, fixed in 2% paraformaldehyde for 15 min on ice, and washed twice in PBS. Cells were then blocked for 1 h in 3% BSA, incubated for 1 h on ice with anti-ErbB3 antibody (1:125; Cat# AF4518, R&D Systems, San Jose, CA, USA), washed twice, and stained for 30 min on ice with Alexa Fluor 488–conjugated secondary antibody (1:250) followed by two final washes. After staining, cells were resuspended in 75 µL PBS and injected into a Moxi GO mini flow cytometer (Orflo Technologies / Precision Cell Systems, Livermore, CA, USA). Stock 2% paraformaldehyde and 3% BSA solutions were prepared by dissolving 2 g PFA or 3 g BSA in 100 mL PBS.

### Biodistribution Assay

Tumor-bearing mice established as described earlier received a single tail vein injection of Her(tcc)P(OH)_2_ or (tcc)P(OH)_2_ alone at 12.5 nmoles Ptcc per injection, then sacrificed and tissue collected at 2-hour (n=3), 4-hour (n=3), and 24-hour (n=3) timepoints. Fluorescence emission (total radiant efficiency) was collected from harvested tissue using an in vivo imaging system (IVIS Spectrum; Perkin Elmer). The total radiant efficiency measured from control organs (non-injected mice) served as background to account for autofluorescence from organs. Quantification of (tcc)P(OH)_2_ per gram of tissue was accomplished by subtracting background from the total radiant efficiency of the harvested organs and extrapolating the values against a standard curve of known titrations of (tcc)P(OH)_2_.

### In vivo studies

Female immunocompetent BALB/c mice (6 weeks old) obtained from Charles River Laboratories received bilateral mammary fat pad implants of 4T1-lucGFP cells (10,000 per implant) and were monitored biweekly via in vivo bioluminescence imaging (BLI) and caliper measurement of primary tumor volumes (depth x width x height) 2-3 times per week until tumors reached ∼100 mm^3^, after which mice were randomized into 8 different treatment groups at N=5 per treatment group. Therapeutic efficacy was evaluated by administering indicated particles or controls via tail vein injection 2 times per week for 5 weeks. Therapeutic dosages of 0.25 mg/kg (based on corrole content) were derived from the minimally effective concentration of corroles used for in vitro therapeutic efficacy. Number and frequency of dosing was based on the calculated particle:cell ratio required for effective response in cell-based assays (# of doses) and the time required for maximum tumor accumulation vs tissue clearance (frequency of dosing), as described previously.(1) Empty particle (corrole-free) dosage was based on protein content of equivalent corrole-containing particles. All reagents were diluted to an injection volume of 0.2 mL with sterile buffer A. Mouse health was monitored regularly by recording weight twice per week. Mouse health was monitored regularly by recording body weight twice per week and by daily visual assessment. Animals were euthanized if tumors exceeded acceptable size limits, appeared ulcerated, or if mice exhibited signs of distress or poor health, in accordance with the approved IACUC protocol (IACUC009356).

### Immunohistology

Slide-mounted tissue specimens were deparaffinized by incubation in a dry oven for 1 hour and then washed with xylene 5 times for 4-minute intervals at a time before rinsing in decreasing ethanol concentrations from 100%-70% (5 times for 4-minute intervals) followed by three 3-minute washes in water rehydrate tissue. Slides were then submerged in antigen retrieval buffer (sodium citrate, pH 6.0) at 95°C for 30-40 minutes and cooled to room temperature before washing 3 times for 5 minutes in DI water. Slides were immersed in permeabilization buffer (PBS w/ Tween-20) for 5 minutes followed by peroxidase inactivation using 3% H2O2 for 30 minutes. Slides were blocked in 1% BSA for 1 hour at room temperature and incubated with primary antibodies (0.2 mL volume) against CD80 (ThermoFisher PA5-85913 1:50), CD8 (Abcam ab25478 1:50) and NK 1.1 (Biolegend 108701 1:50) inside a humified chamber followed by incubation overnight at 4°C. Slides were submerged in PBS w/Tween-20 3 times for 5 minutes to remove any unbound primary antibody followed by exposure to secondary antibodies at 1:400 dilution in blocking buffer (0.2 mL volume) inside humidified chamber for 1 hour in the dark at room temperature. Slides were washed to remove any unbound secondary antibodies and incubated in 0.2 mL volume of True Black lipofuscin autofluorescence quencher (Goldbio cat # TBH-250-1) for 5 minutes followed by rinsing with PBS 3 times for 3 minutes. Slides were counterstained and mounted with 4’6-diamidino-2-phenylindole (DAPI)-containing Prolong Antifade (Thermofisher).

## Data, Materials, and Software Availability

All study data are included in the article and/or SI Appendix.

## ACKNOWLEDGMENTS

Research reported in this publication was supported by grants to L.M.K. from the National Institutes of Health/National Cancer Institute (NIH/NCI; R01 CA258204 and R01 CA270324) and the Department of Defense (DoD; W81XWH-22-1-0953). Research conducted at the California Institute of Technology was supported by the Arnold and Mabel Beckman Foundation. Research conducted at the Technion–Israel Institute of Technology was supported by the Israel Science Foundation (ISF), grant no. 1286/22.

## Author contributions

V.K.S., H.B.G., Z.G., and L.M.K. designed research; V.K.S., N.G.A., S.M., R.H.C., K.I., S.W.K., A.W., R.B. and A.B. performed research; J.A. and R.A. performed MD simulation; V.K.S., N.G.A., S.M., Z.G., and L.M.K. analyzed data; and V.K.S., Z.G., and L.M.K. wrote the paper.

