## Supporting Information for "Next generation protein-corrole bio-assemblies provide effective tumoricidal treatment in a metastatic triple-negative breast cancer model"

**Supplemental Materials**

### Supplemental Movies

Supplemental Movie 1: Molecular dynamics simulation of pentameric HPK2.0 highlighting ligand regions

(https://drive.google.com/file/d/1Ip5L5m8M4a-kOLp6X8krAfKwFdZcc2qZ/view?usp=sharing)

Supplemental Movie 2: Molecular dynamics simulation of pentameric HPK2.0 highlighting cargo loading regions

(https://drive.google.com/file/d/1Ebb9VvvWEuReG-V-KCAzJOuw-fE61Oc9/view?usp=sharing)

### Supplemental Figures

Supplemental Figure S1: HPK1.0 construct, amino acid sequence, and pentameric structure

Supplemental Figure S2: HPK2.0 construct, amino acid sequence, and pentameric structure

Supplemental Figure S3: MD simulation snapshot of HPK1.0 showing K10 tail positioning

Supplemental Figure S4: UV/Vis spectra of S2Ga, (tcc)Ga, and (tcc)P$(OH)_2$

Supplemental Figure S5: ITC analysis of (tcc)Ga binding to HPK2.0

Supplemental Figure S6: Photoluminescence spectra of free and HPK2.0-bound (tcc)Ga

Supplemental Figure S7: Visual assessment of corrole–HPK2.0 bio-assemblies

Supplemental Figure S8: DLS size distributions and long-term stability analysis

Supplemental Figure S9: Evaluation of HPK2.0–corrole assemblies on cell line growth.

Supplemental Figure S10: In vivo tumor growth inhibition by (tcc)P$(OH)_2$ and S2Ga bio-assemblies

Supplemental Figure S11: Kaplan–Meier survival analysis

Supplemental Figure S12: Body weight monitoring during treatment

Supplemental Figure S13: Tumor-resident immune cell analysis following treatment

### Supplemental Tables

Supplemental Table S1: $IC_{50}$ values in 4T1 cells under dark and light conditions

Supplemental Table S2: $IC_{50}$ values in NIH-3T3 cells under dark and light conditions

**Supplemental Movie 1.** Molecular dynamics simulation of pentameric HPK2.0 highlighting ligand regions. **Movie S1** shows ribbon structure of pentameric HPK2.0 with its 5 ligand regions (that engage the HER3 receptor) delineated by gold color. Movie stills of key sequential time points are shown below (ns, nanoseconds).

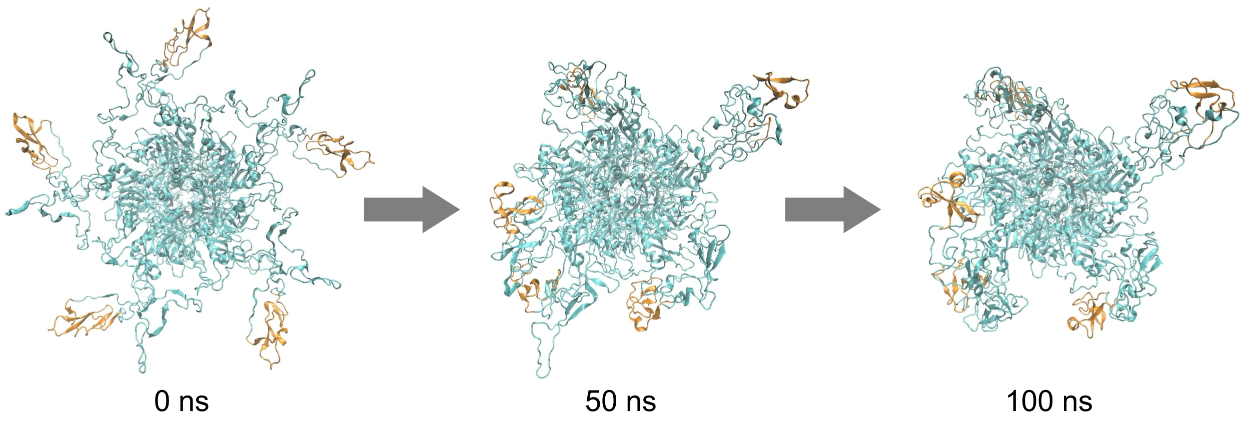


**Supplemental Movie 2.** Molecular dynamics simulation of pentameric HPK2.0 highlighting cargo loading regions. **Movie S2** shows ribbon structure of pentameric HPK2.0 with its 5 carboxy-terminal decalysine tails delineated by red color. Movie stills of key sequential time points are shown below (ns, nanoseconds).

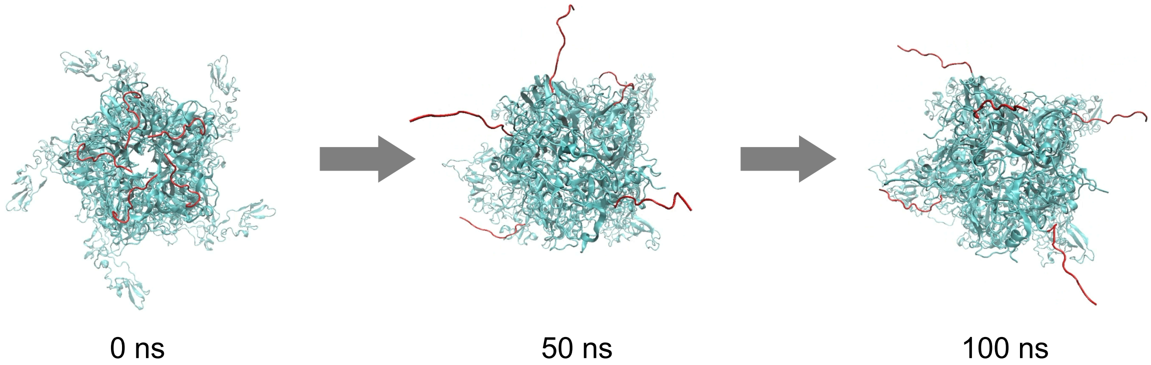

A

**Affinity tag: 1-36**
**Targeting ligand: Her (Ig-like, 37-122; EGF-like, 123-243)**
**Linker: 244-251**
**PB: 252-822**
**K10: 823-832**

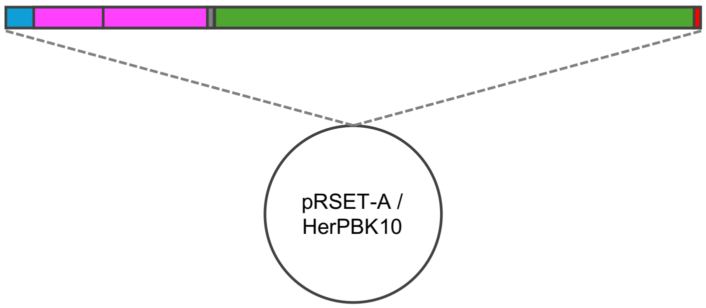


B

**HerPBK10** **(HPK1.0):**

MRGSHHHHHHGMASMTGGQQMGRDLYDDDDKDRWGSELLPPQLKEMKSQESAAGS
KLVLRCETSSEYSSLRFKWFKNGNELNRKNKPQNIKIQKKPGKSELRINKASLAD
SGEYMCKVISKLGNDSASANIAIVESNEIITGMPASTEGAYVSSESPIRISVSTE
GANTSSSTSTSTTGTSHLVKCAEKEKTFCVNGGECFMVKDLSNPSRYLCKCQPGF
TGARCTENVPMKVQNQEKAEELYGGSGGSRSMRRAAMYEEGPPPSYESVVSAAPV
AAALGSPFDAPLDPPFVPPRYLRPTGGRNSIRYSELAPLFDTTRVYLVDNKSTDV
ASLNYQNDHSNFLTTVIQNNDYSPGEASTQTINLDDRSHWGGDLKTILHTNMPNV
NEFMFTNKFKARVMVSRLPTKDNQVELKYEWVEFTLPEGNYSETMTIDLMNNAIV
EHYLKVGRQNGVLESDIGVKFDTRNFRLGFDPVTGLVMPGVYTNEAFHPDIILLP
GCGVDFTHSRLSNLLGIRKRQPFQEGFRITYDDLEGGNIPALLDVDAYQASLKDD
TEQGGGGAGGSNSSGSGAEENSNAAAAAMQPVEDMNDHAIRGDTFATRAEEKRAE
AEAAAEAAAPAAQPEVEKPQKKPVIKPLTEDSKKRSYNLISNDSTFTQYRSWYLA
YNYGDPQTGIRSWTLLCTPDVTCGSEQVYWSLPDMMQDPVTFRSTRQISNFPVVG
AELLPVHSKSFYNDQAVYSQLIRQFTSLTHVFNRFPENQILARPPAPTITTVSEN
VPALTDHGTLPLRNSIGGVQRVTITDARRRTCPYVYKALGIVSPRVLSSRTFKKK
KKKKKKK

C

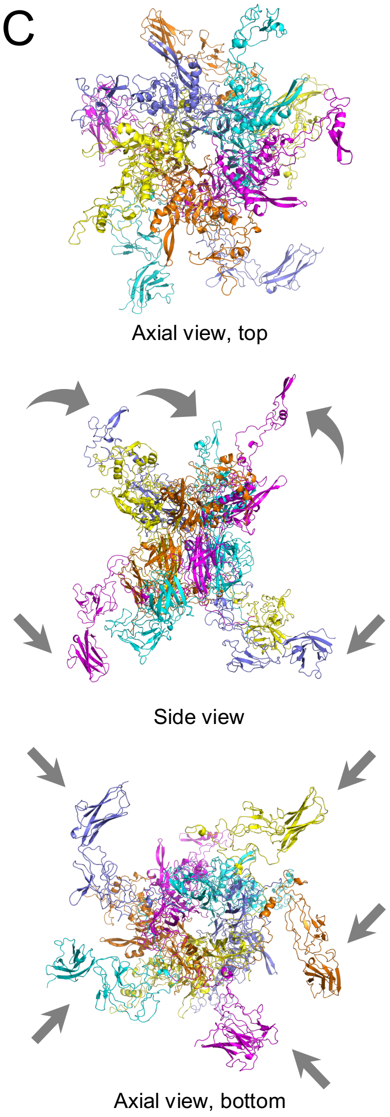


**Supplemental Figure S1.** HPK1.0. **A**, Linear depiction of HPK1.0 gene sequence. Segments encoding key functional domains are color coded. Numbering corresponds to amino acid sequence from amino [N] to carboxy [C] terminus. **B**, Amino acid sequence of HPK1.0 with key functional domains color coded to correspond to the regions depicted in ***A***. **C**, Ribbon structure of HPK1.0 pentameric capsomere generated by computational modeling. Each monomer is delineated by a different color. Straight arrows point to targeting ligands; curved arrows point to the RGD loops in the penton base domains.

A

**His: 1-36**
**PB1:37-321**
**L1: 322-337 (double linker)**
**EGF: 338-449**
**L2: 450-465 (double linker)**
**PB2: 466-698**
**L3: 699-706 (single linker)**
**K10: 707-716**

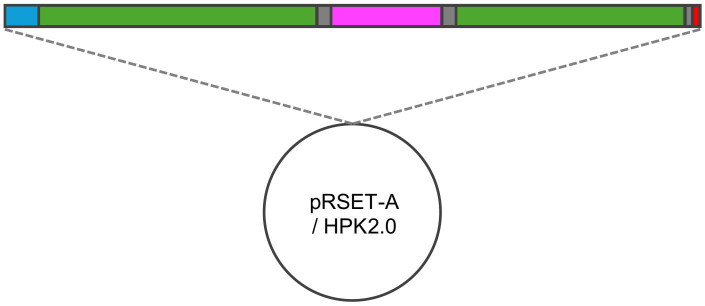


B

**PB(Δ50ΔRGD)Her(ΔIgG)K10+Extra Linkers** **<u>(HPK 2.0)</u>**

```
MRGSHHHHHHGMASMTGGQQMGRDLYDDDDKDRWGSGRNSIRYSELAPLFDTT
RVYLVDNKSTDVASLNYQNDHSNFLTTVIQNNDYSPGEASTQTINLDDRSHWG
GDLKTILHTNMPNVNEFMFTNKFKARVMVSRLPTKDNQVELKYEWVEFTLPEG
NYSETMTIDLMNNAIVEHYLKVGRQNGVLESDIGVKFDTRNFRLGFDPVTGLV
MPGVYTNEAFHPDIILLPGCGVDFTHSRLSNLLGIRKRQPFQEGFRITYDDLE
GGNIPALLDVDAYQASLKDDTEQGGGGAGGSNSSGSGAEENSNAAAAAMQPVE
DMNGGSGGSRSGGSGGSRSAIVESNEIITGMPASTEGAYVSSESPIRISVSTE
GANTSSSTSTSTTGTSHLVKCAEKEKTFCVNGGECFMVKDLSNPSRYLCKCQP
GFTGARCTENVPMKVQNQEKAEELYGGSGGSRSGGSGGSRSDHATIFATRAEE
KRAEAEAAAEAAAPAAQPEVEKPQKKPVIKPLTEDSKKRSYNLISNDSTFTQY
RSWYLAYNYGDPQTGIRSWTLLCTPDVTCGSEQVYWSLPDMMQDPVTFRSTRQ
ISNFPVVGAELLPVHSKSFYNDQAVYSQLIRQFTSLTHVFNRFPENQILARPP
APTITTVSENVPALTDHGTLPLRNSIGGVQRVTITDARRRTCPYVYKALGIVS
PRVLSSRTFGGSGGSRSKKKKKKKKKK
```

C

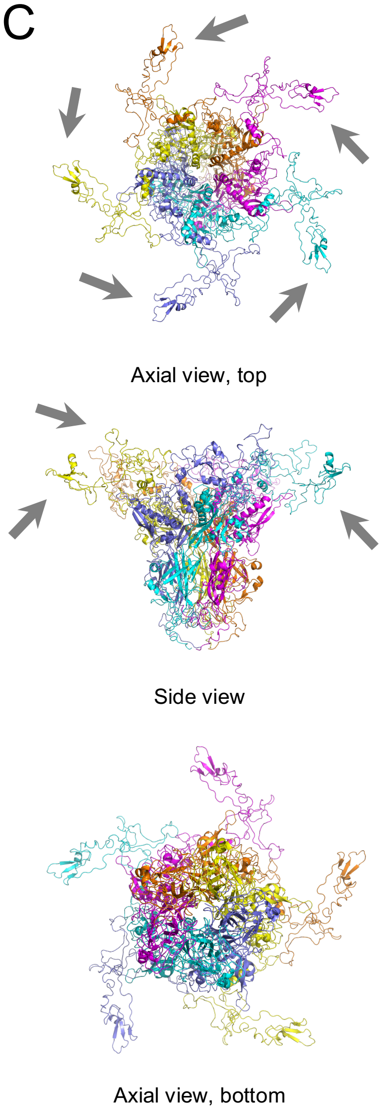


**Supplemental Figure S2.** HPK2.0. **A**, Linear depiction of HPK2.0 gene sequence. Segments encoding key functional domains are color coded. Numbering corresponds to amino acid sequence from amino [N] to carboxy [C] terminus. **B**, Amino acid sequence of HPK2.0 with key functional domains color coded to correspond to the regions depicted in ***A***. **C**, Ribbon structure of HPK2.0 pentameric capsomere generated by computational modeling. Each monomer is delineated by a different color. Arrows point to targeting ligands.

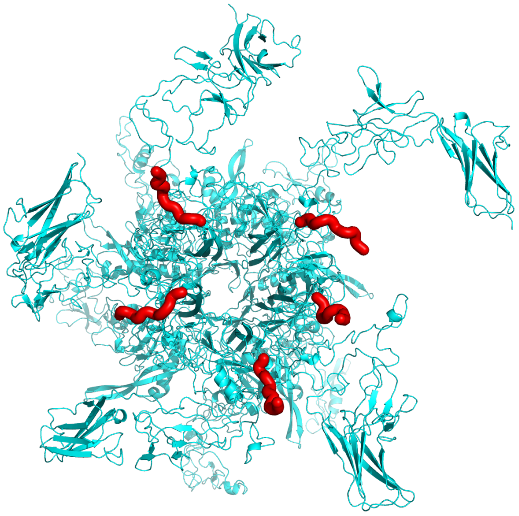

**Supplemental Figure S3**. Computational model of HPK1.0 pentamer shown in ribbon structure form, highlighting the 5 K10 tails in red.

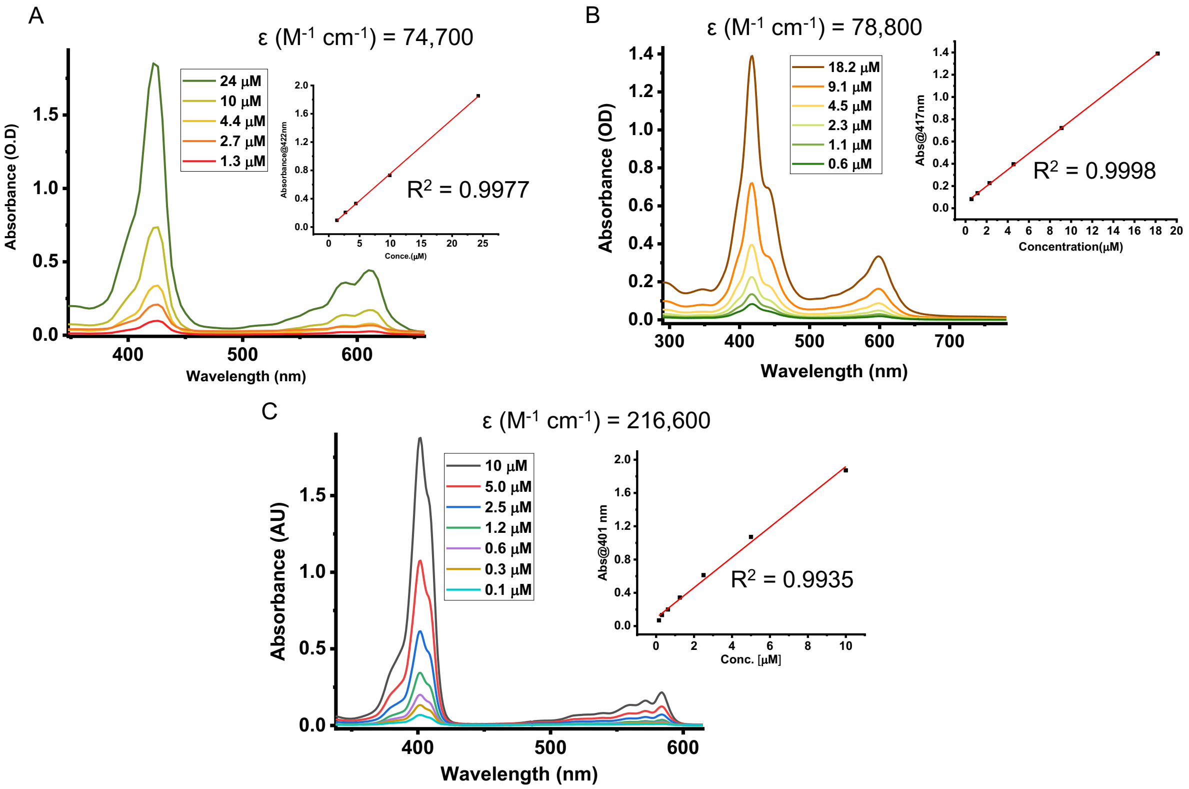


**Supplemental Figure S4.** UV/vis spectra of (**A**) S2Ga, (**B**) (tcc)Ga, and (**C**) (tcc)$P(OH)_2$ in 100 mM PBS at pH 7.4. The inset shows the absorbance versus concentration plots. The linear relationship demonstrates adherence to the Beer-Lambert Law. The slopes correspond to the molar extinction coefficients (ε) of the compounds.

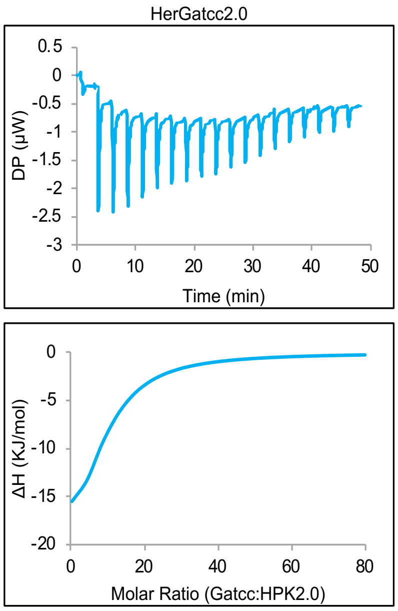


| N (sites) | KD (M) | ΔH (kJ/mol) | ΔG (kJ/mol) | -TΔS (kJ/mol) |
|---|---|---|---|---|
| 8 | 21.1e-6 | -27.3 | -26.7 | 0.60 |

**Supplemental Figure S5.** Isothermal Titration Calorimetry (ITC) analysis of (tcc)Ga binding to HPK2.0. Below are the thermodynamic parameters summarized in tabular form for the formation of bio-assemblies.

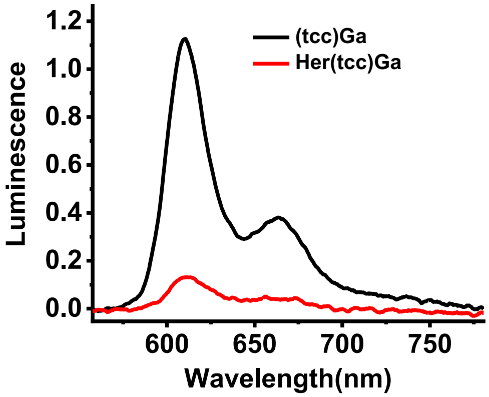


**Supplemental Figure S6.** Photoluminescence spectra of unbound (black traces) and HPK2.0-bound (red traces) (tcc)Ga.

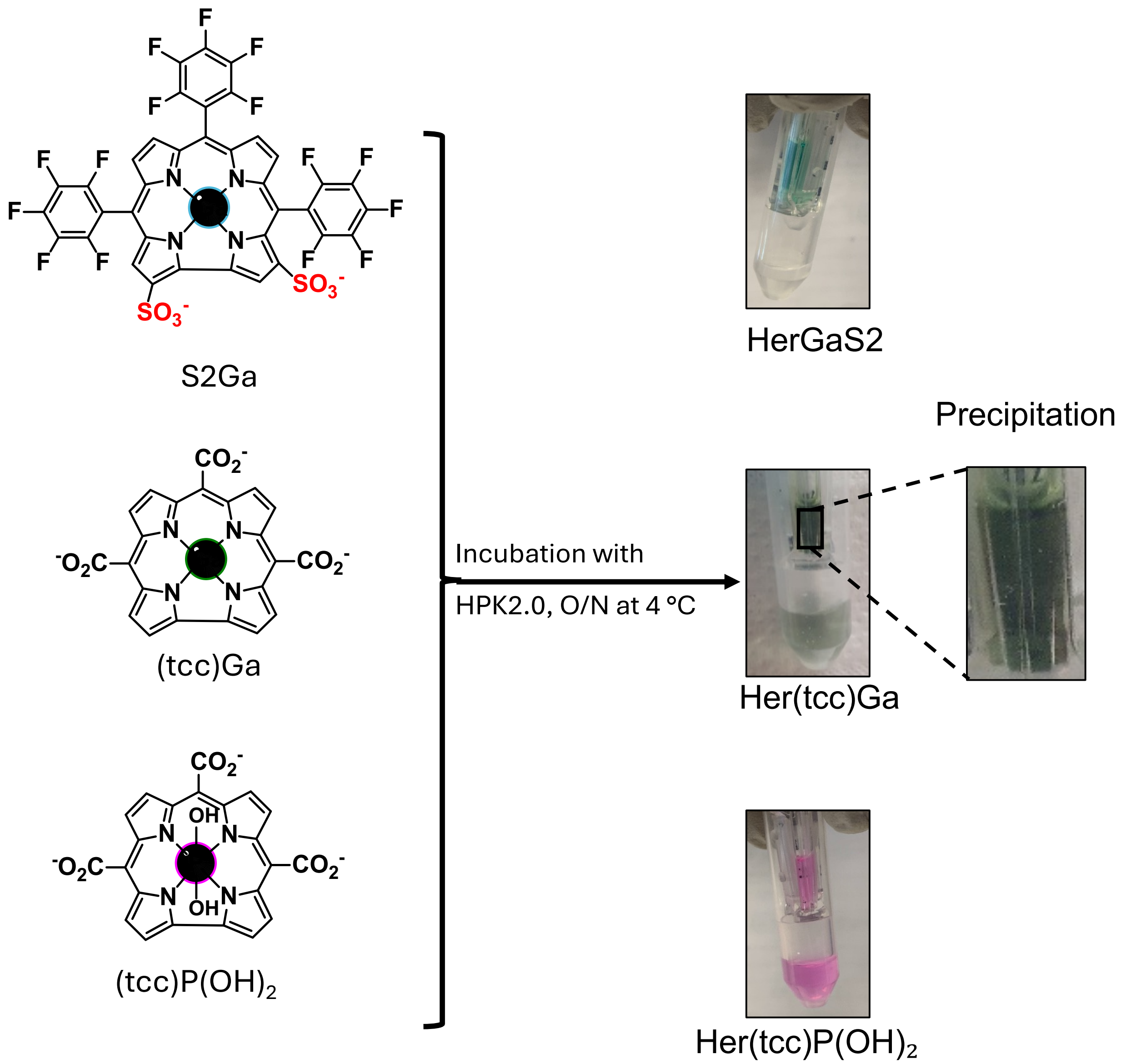


**Supplemental Figure S7.** Visual outcomes of corrole–HPK2.0 assembly reactions after indicated corroles [$S_2Ga$, (tcc)Ga, (tcc)$P(OH)_2$] were each incubated with HPK2.0 overnight at 4 °C, followed by ultrafiltration of each mixture. Photos (right) capture both retentates and filtrates of each mixture with the distribution of each corrole traceable based on its corresponding pigment (and confirmed by optical spectroscopy).

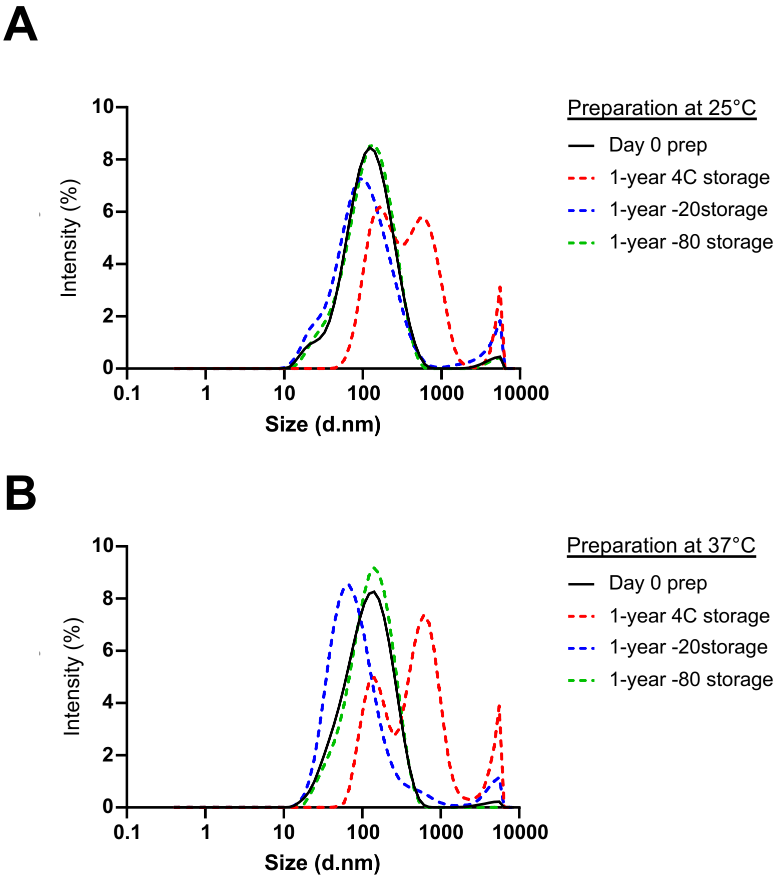


**Supplemental Figure S8.** DLS spectra (measured at 25 °C) of (tcc)P$(OH)_2$ bio-assemblies prepared at (**A**) 25 °C , or (**B**) 37 °C . Solid lines represent size distributions on the day of assembly preparation; dashed lines show distributions after 1 year of storage at 4 °C, –20 °C, or –80 °C.

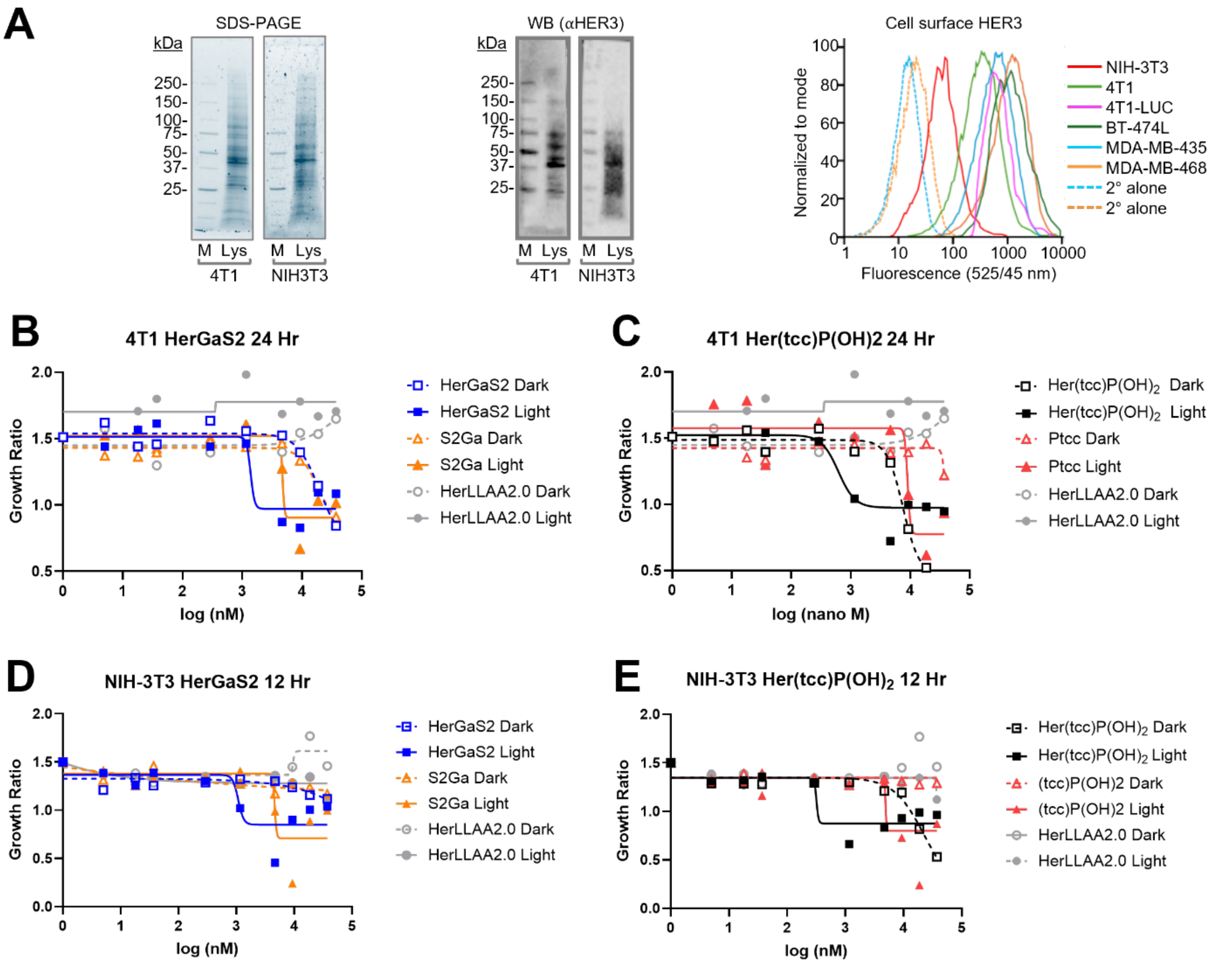


**Supplemental Figure S9.** Evaluation of HPK2.0–corrole assemblies on cell line growth. **A**, Cell line HER3 densities. SDS-PAGE (left) and immunoblots (middle) of indicated cell lysates probed for HER3. Right, relative cell surface HER3 on indicated cell lines. Dotted lines match corresponding controls (MDA-MB-435, MDA-MB-468) treated with secondary antibody alone. **B-E**, Evaluation of cytotoxicity -/+ photo-activation. Graphs summarize growth inhibition curves of mouse TNBC (4T1) (**B-C**) or fibroblasts (NIH-3T3) (**D-E**) exposed to the indicated bio-assemblies under dark conditions (open symbols) or with brief photoactivation (closed symbols) (details in the *Methods*).

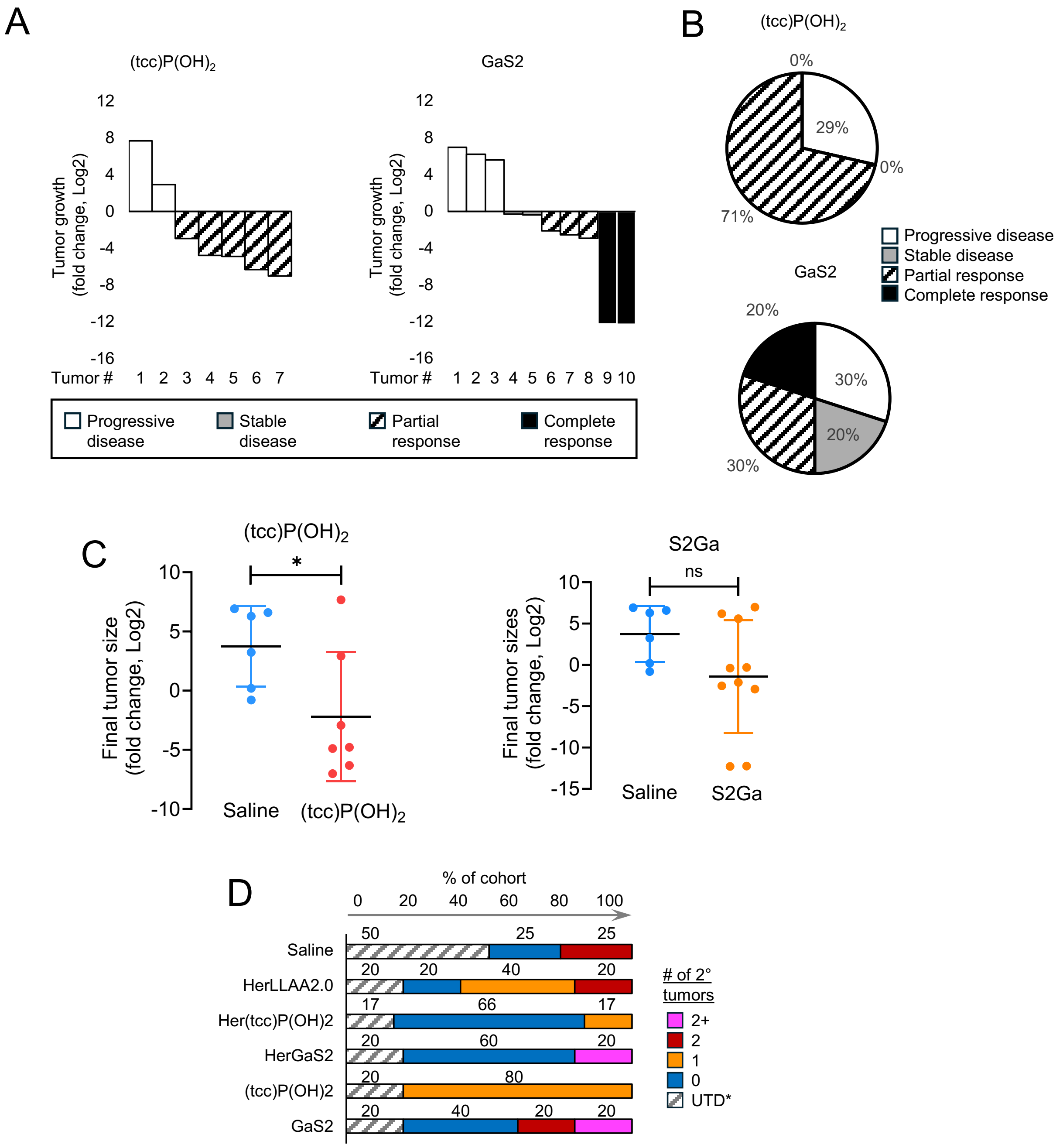


**Supplemental Figure S10.** Effect of corrole bio-assemblies on tumor growth *in vivo*. **A**, Waterfall plots showing $\log_2$ fold change in primary tumor growth for individual tumors following treatment with $(tcc)P(OH)_2$ or GaS2 at a dose of 0.22 mg/kg. Tumors are classified according to RECIST-adapted response criteria: progressive disease, stable disease, partial response, or complete response. **B,** Pie charts summarizing the distribution of RECIST-adapted therapeutic response categories for each treatment cohort. Percentages indicate the fraction of tumors in each category. **C,** Scatter plot of $\log_2$ fold change in final primary tumor sizes for individual tumors across treatment groups. Horizontal lines indicate group means. Statistical comparisons are shown as indicated (ns, not significant; $^*p < 0.05$). **D,** Distribution of the number of secondary tumors per mouse across treatment cohorts. *Cohort sizes ranged from n = 4–6 mice per treatment group.*

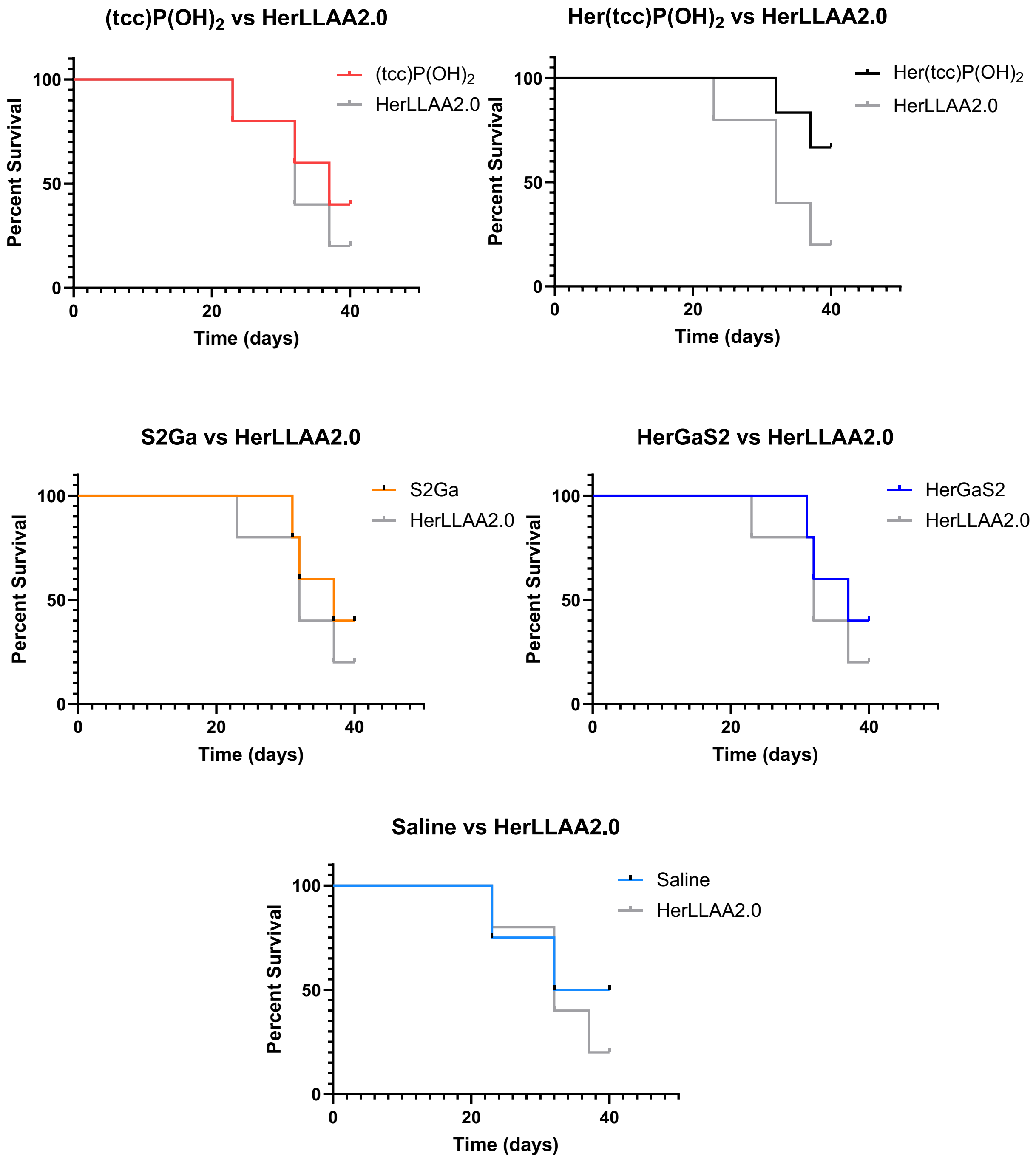


**Supplemental Figure S11.** Kaplan–Meier survival analysis of mice treated with nano-assemblies, corrole alone, saline (control), or HerLLAA2.0 (control). Each panel shows survival curves for individual treatment groups compared against controls. Mice were monitored daily for survival following treatment. Curves represent the percentage of surviving animals over time (days).

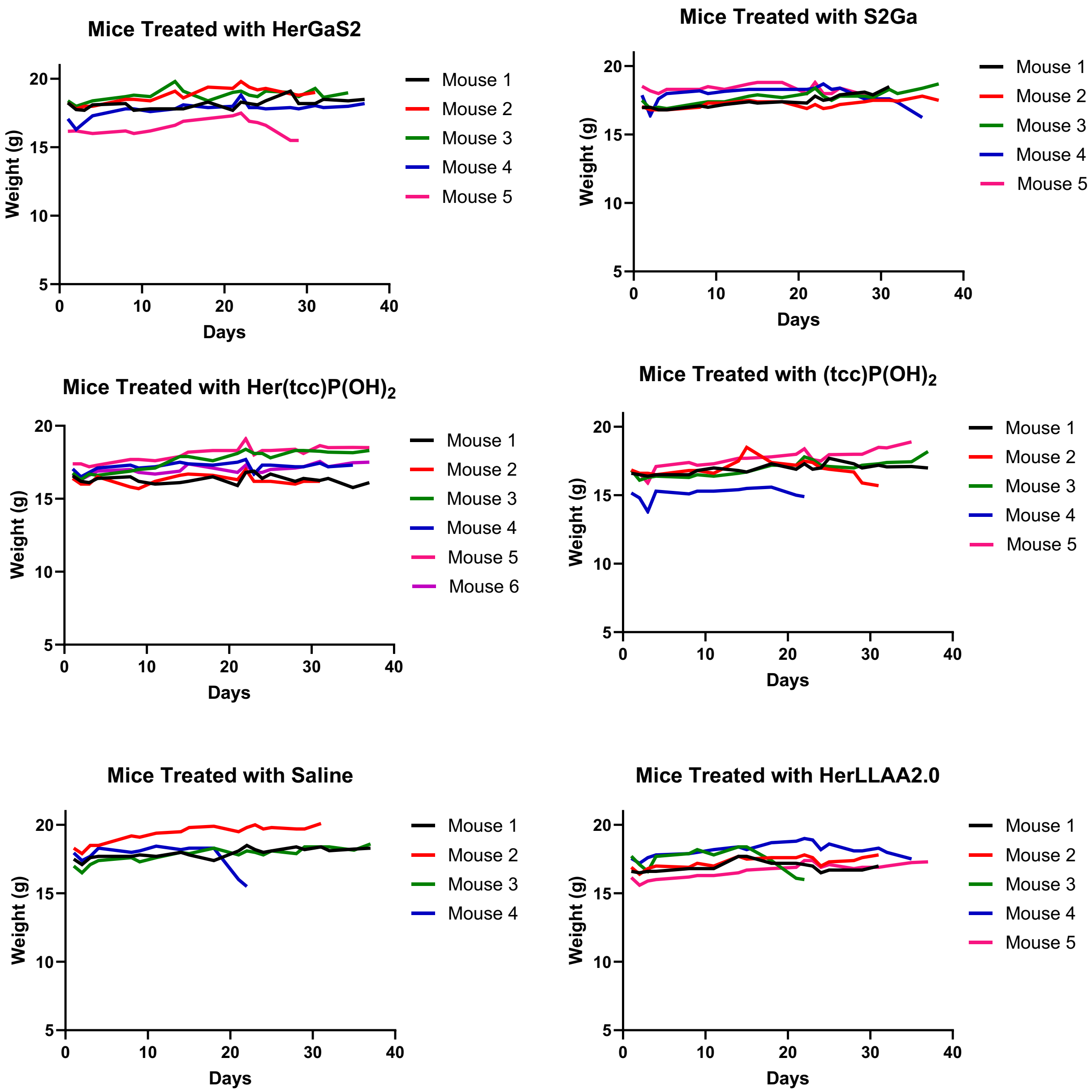


**Supplemental Figure S12.** Body weight of mice treated with different drugs over time. Each panel shows body weight (grams) measured over time for separate experimental groups. Colored lines correspond to individual mice that each received a particular drug treatment. Body weights were recorded twice per week throughout the study.

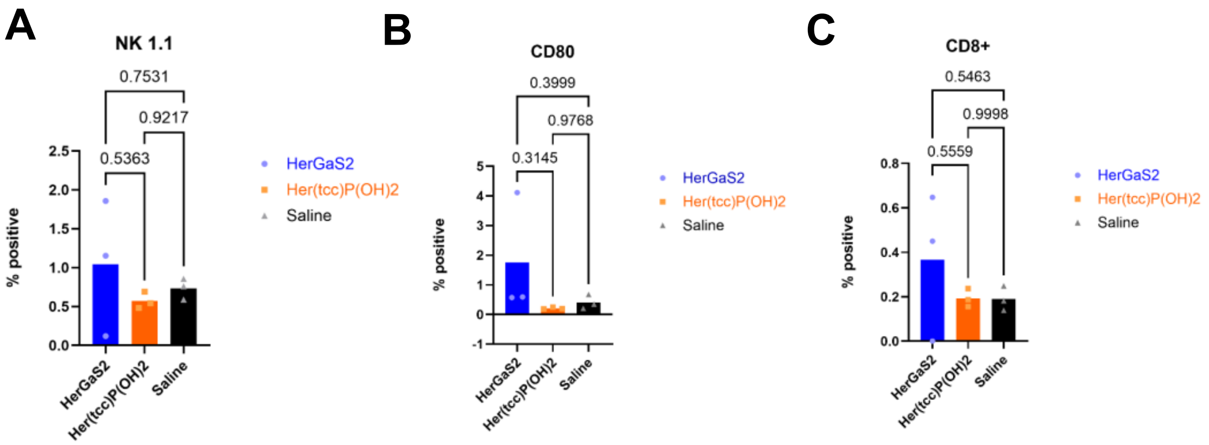


**Supplemental Figure S13.** Tumor-resident immune cell populations following HPK2.0–corrole therapy. Orthotopic HER3$^{+}$ 4T1 tumors were harvested from BALB/c mice after systemic treatment with HerGaS2, Her(tcc)P(OH)$_2$, or saline (vehicle control). Immune cell infiltration was quantified by immunofluorescence staining for: (**A**) NK1.1$^{+}$ natural killer cells, (**B**) CD80$^{+}$ activated antigen-presenting cells, and (**C**) CD8$^{+}$ cytotoxic T cells. Immune cell levels were low across all treatment groups, and no significant differences were observed, indicating that HPK2.0–corrole assemblies do not generate detectable immune activation within the tumor microenvironment under therapeutic conditions. Data represent individual mice (n = 3/group); bars show mean ± SEM. *P*-values (one-way ANOVA with Tukey's multiple-comparisons test) are indicated above comparison brackets.

| Name | IC50 on 4T1 (nM) | IC50 on 4T1 (μM) |
|---|---|---|
| HerGaS2 Dark | 19995 | 20 |
| HerGaS2 Light | 1380 | 1.4 |
| S2Ga Dark | 18353 | 18.4 |
| S2Ga Light | 4723 | 4.7 |
| Her(tcc)P(OH)$_2$ Dark | 7563 | 7.6 |
| Her(tcc)P(OH)$_2$ Light | 619 | 0.6 |
| (tcc)P(OH)$_2$ Dark | 101514 | 101.5 |
| (tcc)P(OH)$_2$ Light | 9032 | 9 |
| HerLLAA2.0 Dark | 38910 | 38.9 |
| HerLLAA2.0 Light | Unstable | Unstable |

**Supplemental Table 1**. Comparison of cytotoxic potency ($IC_{50}$) of all formulations in 4T1 cells under dark and light conditions. $IC_{50}$ values are reported in both nanomolar (nM) and micromolar (μM) units. Several formulations show enhanced potency upon light exposure, with Her(tcc)P(OH)$_2$ exhibiting the lowest $IC_{50}$ under light conditions. Free corroles displayed minimal activity, and HerLLAA2.0 showed instability under light exposure. Data represent mean values from n = 3 biological replicates per concentration. $IC_{50}$ values were calculated using nonlinear regression with a four-parameter logistic model.

| Name | IC50 on NIH-3T3 (nM) | IC50 on NIH-3T3 (µM) |
|---|---|---|
| HerGaS2 Dark | 3.2195E+13 | 32195035690 |
| HerGaS2 Light | 1075 | 1.075 |
| S2Ga Dark | 1.058E-24 | 1.058E-27 |
| S2Ga Light | 4593 | 4.593 |
| Her(tcc)P$(OH)_2$ Dark | 22445 | 22.445 |
| Her(tcc)P$(OH)_2$ Light | 318.6 | 0.3186 |
| (tcc)P$(OH)_2$ Dark | Unstable | Unstable |
| (tcc)P$(OH)_2$ Light | 4892 | 4.892 |
| HerLLAA2.0 Dark | Unstable | Unstable |
| HerLLAA2.0 Light | 46593 | 46.593 |

**Supplemental Table 2**. Comparison of cytotoxic potency ($IC_{50}$) of all formulations in NIH-3T3 cells under dark and light conditions. $IC_{50}$ values are reported in both nanomolar (nM) and micromolar (µM) units. Most formulations exhibited limited cytotoxicity in the dark, while select formulations showed increased activity upon light exposure. Several conditions resulted in unstable or non-quantifiable $IC_{50}$ values. Data represent mean values from n = 3 biological replicates per concentration. $IC_{50}$ values were determined by nonlinear regression using a four-parameter logistic curve fit.
